# Enhancing Task fMRI Preprocessing via Individualized Model-Based Filtering of Intrinsic Activity Dynamics

**DOI:** 10.1101/2020.12.10.420273

**Authors:** Matthew F. Singh, Anxu Wang, Michael Cole, ShiNung Ching, Todd S. Braver

## Abstract

Brain responses recorded during fMRI are thought to reflect both rapid, stimulus-evoked activity and the propagation of spontaneous activity through brain networks. In the current work we describe a method to improve the estimation of task-evoked brain activity by first “filtering-out” the intrinsic propagation of pre-event activity from the BOLD signal. We do so using Mesoscale Individualized NeuroDynamic (MINDy; [1]) models built from individualized resting-state data (MINDy-based Filtering). After filtering, time-series are analyzed using conventional techniques. Results demonstrate that this simple operation significantly improves the statistical power and temporal precision of estimated group-level effects. Moreover, use of MINDy-based filtering increased the similarity of neural activation profiles and prediction of individual differences in behavior across tasks measuring the same construct (cognitive control).Thus, by subtracting the propagation of previous activity, we obtain better estimates of task-related neural activity.

## 1. Introduction

Task-related analyses in fMRI typically involve statistical general linear models (GLMs) which seek to identify the amplitude and/or mean timecourse of (BOLD) evoked-responses after removing nuisance covariates. These approaches have proven statistically powerful and characterize much of the current literature regarding task-induced activation in group-level fMRI analyses. However, over the past two decades, improvements in fMRI acquisitions and the rise of resting-state connectomics ([2]) have given rise to a new literature concerning variability within brain activation across trials, individuals, and/or contexts. Understanding such variability is key to precision neuroscience initiatives, as these studies have the potential to uncover new neural mechanisms and generate stronger brain-behavior linkages at the level of individuals ([3], [4], [5]).

Previous studies in this domain have generated two key findings relevant to the current study: 1) individual differences in intrinsic brain networks predict corresponding differences in BOLD responses ([6], [7], [8], [9]) and 2) the BOLD signal elicited by a stimulus is dependent upon the previous pattern of brain activity ([10]), including spontaneous fluctuations ([11]). We use the term “brain activity” in the latter case to indicate that this history dependence is thought to be neural, rather than solely reflecting potential nonlinearity in the hemodynamic coupling. The first set of findings indicate that inter-subject variability in brain responses may be due to the “flow” ([8]) of evoked activity through subject-specific connectomes. The second set of findings suggest that evoked responses are history-dependent (i.e. reflects underlying dynamics). Thus, the neural activity associated with BOLD is increasingly considered as a nonlinear dynamical system—one in which the spatiotemporal response to an input depends upon its current state, and further, is determined by a set of rules that dictate its temporal evolution ([12]). These dynamical “rules” are a function of subject-specific connectivity and the specific properties local to each brain region ([13], [14]). The manifestation of these dynamics (i.e. trial-to-trial variability in BOLD) are thought to be neural and cognitively-relevant as they predict within-subject behavioral variation ([15]).

This framework contrasts both with current statistical approaches, which treat the neural activity as a noisy autoregressive signal (most GLMs), and with Dynamic Causal Modeling (DCM) approaches, which treat the brain as a linear system (although see [16]). In the current work, we propose a new technique for modeling intrinsic brain dynamics and their contribution to task-evoked activation patterns. This approach leverages MINDy models ([1]) fit to resting-state data for each subject. These models are akin to an abstracted neural mass model containing hundreds of different regions (parcels) spanning the whole brain. Regions interact nonlinearly via a signed, directed connectivity matrix and integrate inputs over time (i.e. form a nonlinear dynamical system). The BOLD signal is modeled via region-specific hemodynamic models, and all parameters (neural and hemodynamic) are directly estimated from each subject’s resting-state scans (a process which takes 1-3 minutes). In prior work ([1], [17]), we have established that MINDy models/parameters are robust, reliable, and predictive ([1]). In the current work, we use these models to estimate intrinsic brain dynamics (i.e. predictions based upon resting-state MINDy models) and subtract them from the observed BOLD, a process which we term MINDy-based Filtering. This procedure more sensitively identifies individual differences, and enhances the temporal precision and statistical power in identifying task effects. We also obtain stronger brain-behavior linkages and greater similarity in the activation profiles of different tasks that index a common cognitive construct (cognitive control demand).

### 1.1. Filtering Intrinsic Dynamics

The current approach rests upon the ability to model the flow of neural activity between brain areas, as identified via models fit to resting-state brain activity. However, rather than seeking to describe the flow of task-related neural activity (e.g. [8]), our approach acts to censor, or computationally estimate and remove, the flow of task-unrelated (pre-event) activity. To be clear, we perform this operation at every time point and use the whole timeseries for analyses. No information regarding task timing is used in our filter (Fig. 1A). However we use the notion of “events” to provide an intuitive motivation for our approach (conversely each timepoint could be considered an “event”). Likewise, our approach does not require an event-related design (see SI 7.5 for block-related analyses). At each time point, the measured neural activity is considered a combination of task-evoked effects manifest over fast time scales and the propagation of brain activity emerging at previous time points. By subtracting the modeled propagation of previously-triggered (e.g. pre-event) activity, we aim to better isolate the influence of each event (time-point).

**Fig 1.**
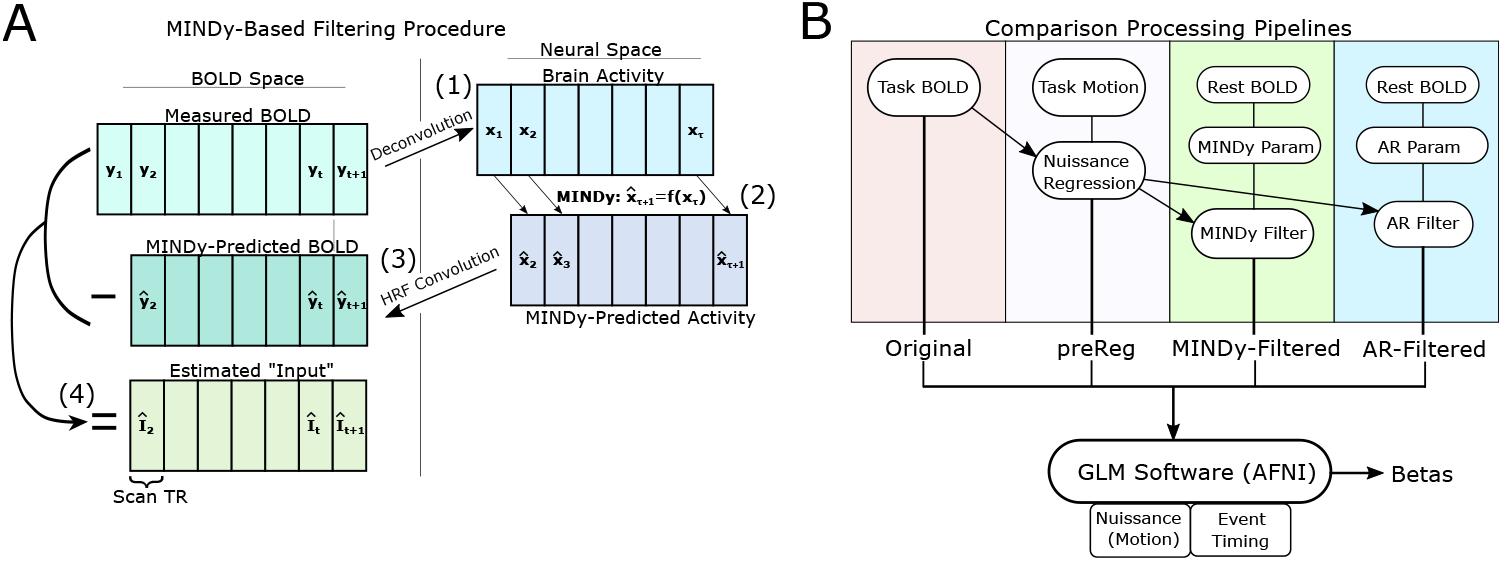
Filtering and control pipelines. A) MINDy-based Filtering procedure. 1) Latent neural activity is estimated from the BOLD signal. 2) One-step predictions for latent neural activity are made with MINDy and 3) convolved into one-step BOLD predictions. 4) Filtered “input”/residual timeseries are the difference of measured and predicted BOLD (we abbreviate *h \** as *Î*). B) Analysis pipelines. Modeling pipelines require data to be pre-processed (nuissance regressed) before model-based filtering. The preReg pipeline controls for this step by performing identical pre-processing before GLM analyses. Parameters for MINDy and autoregressive models are estimated from resting-state data. Autoregressive models (AR) are used to test whether effects are due to local signal-processing features (i.e. MINDy similar to AR) vs. exploit brain connectivity (MINDy better than AR). Although we chose AFNI to perform GLM analyses, MINDy-based Filtering is compatible with any analysis software as filtered timeseries are analyzed in the conventional manner.

Our approach is conceptually-similar to a previous study by Fox and colleagues ([11],[18]) which suggested that estimated task-effects could be improved by subtracting spontaneous activity. They demonstrated this possibility in a motor task by subtracting the recorded BOLD in contralateral motor cortex from the task-implicated motor hemisphere. However, the Fox et al. approach ([11],[18]) has not been applied more broadly, since it requires identifying region pairs which are strongly correlated at rest, but only one of which is recruited during task. This dissociation is key as it enabled Fox and colleagues ([11]) to measure intrinsic brain activity (via the contralateral cortex) separately from task-evoked activity in the other hemisphere. By contrast, the current literature overwhelmingly suggests that, for most brain regions and networks, coactivation during resting-state fMRI predicts coactivation during task (e.g. [6], [8], [7]).

By contrast, we propose to filter out the intrinsic component of brain activity using model-based predictions. We predict brain activation at each time-point by applying MINDy models derived from resting-state activity ([1],[17]) to the previous time-step (i.e. 1-step forward predictions) and subtract these predictions to better identify task-evoked changes. Thus, we better isolate event-related brain changes by filtering out the propagation of pre-event activity. As mentioned previously, we use the notion of task “events” to provide an intuitive understanding of why our approach improves fMRI analyses. Our filter does not utilize any prior information regarding task structure (events) and is compatible with any task design (not just event-related designs; see Fig. 1B).

### 1.2. Previous Approaches using DCM

Dynamic Causal Modeling (DCM), by contrast, incorporates the temporal evolution of brain activity and thus can consider the propagation of neural activity through brain networks. Each DCM contains an effective connectivity matrix and a set of extrinsic inputs that describe how task events impinge upon each node of the network ([19]). Many implementations also contain region-specific hemodynamic models and/or an interaction between task events and effective connectivity (i.e., the effective connectivity is parameterized by task events). Although the original DCM models were strongly limited in size, modern implementations ([20], [21]) can consider a much larger number of brain regions (although the computation cost still remains considerable; [20], [1]). However, the DCM methodology also presents several constraints which limit its application. Estimating a DCM model requires pre-specifying the time-series of task effects. This assumption precludes analyses which explore the temporal dynamics of task effects such as Finite Impulse Response (FIR) modeling or nuanced task GLMs, such as those featuring nuisance regressors (e.g. motion). In addition, all DCM implementations that support whole-brain models (i.e., more than a few regions; [20]) are dependent upon the assumption of stationary linear dynamics ([1]).

### 1.3. Filtering Instead of Parameterizing

In the current work, we aim to strike a balance between the mechanistic inferences made by DCM and the flexibility of standard analysis techniques. To do so, we generate dynamical systems models of the brain and neurovasculature (as is done in DCM). However, our approach differs substantially from DCM in how we build and utilize these models. Instead of fitting models of the brain and tasks, we propose to fit dynamic models to independent resting-state data for each subject. We then use these models to generate a mathematical filter for each subject that removes, or “partials out”, the effects of intrinsic dynamics from BOLD timeseries. The approach uses no information regarding task events and thus functions as a preprocessing step, as opposed to explicitly modeling task events. This feature is advantageous, as the proposed techniques can be inserted into any data preprocessing pipeline with minimal effort, provided that sufficient amount of resting state data (e.g. *>*15 minutes [1]) has been collected to build MINDy models.

## 2. Approach

In our approach we predict future BOLD measurements, while modeling biological activity at the neural (i.e., deconvolved) level. Generative models are parameterized according to resting-state data. The MINDy-Filtered data is defined by the difference between measured and model-predicted BOLD. Our procedure thus contains two stages: (1) parameterizing resting-state MINDy models; and (2) using these models to perform MINDy-based Filtering. We begin by reviewing the resting-state MINDy model.

### 2.1. Resting-State MINDy Modeling

The MINDy model ([1],[17]) is a phenomenological extension of neural-mass type models which operates at timescales commensurate with fMRI. Like neural-mass models, MINDy models contain three components: a signed, directed weight matrix of estimated effective connectivities (*W*), a sigmoidal transfer function (*ψ*) which relates local activation to the strength of outward signaling, and the region-specific decay rate (time-constant) *D* which describes how quickly a stimulated region will return to baseline levels of activity. MINDy models operate at two time-frames: the time-frame of neural activity (denoted *τ*) and the time-frame of BOLD measurements (denoted *t*) which we assume are linked by region-specific hemodynamic-response-function *h_β_*. The resting-state neural activity (*x_τ_*) evolves according to the discrete-time dynamical system:

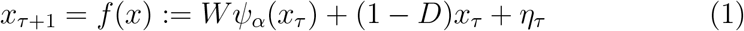

with process noise *η_t_* assumed uncorrelated between parcels. The transfer function *ψ* is parameterized by the curvature vector *α* which dictates regional-differences in the shape of *ψ*:

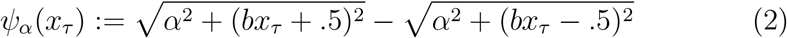

with *b* = 20*/*3 a fixed, global hyperparameter. These neural equations are linked to the observed BOLD measurements via the convolutional HRF model. We model HRFs using a parameterized version of the canonical double-gamma model with vector-valued parameters *β*_1_*, β*_2_:

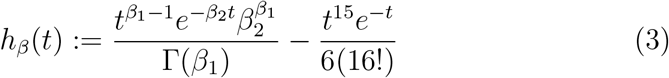

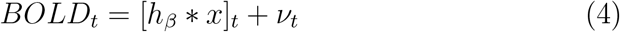

MINDy quickly and simultaneously solves for *W, α, D*, and *β* using a unique, regularized optimization method ([1],[17]). Neural states are inverted from BOLD using the Wiener deconvolution ([22]). Denoting complex-conjugation by *z^∗^*, the Fourier-transform by 𝓕 and the Wiener NSR parameter *ε* = .002 (see SI Sec. 7.4), we define the Wiener HRF-deconvolution (𝓗^+^) as:

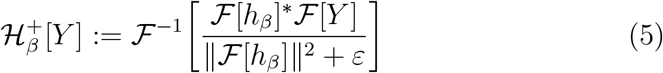

All multiplications/divisions in the above equation are understood to be element-wise. We similarly implement convolution using the Fourier transform (by the Convolution Theorem: 𝓕[*x ∗ y*] = 𝓕[*x*]𝓕[*y*]):

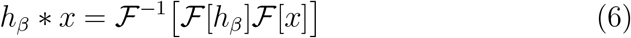

Thus, the combined MINDy model for resting-state (excluding noise) is:

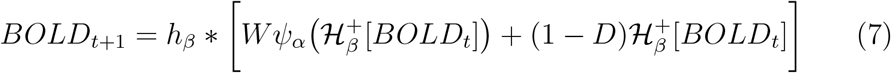

Since the exact convolution and deconvolution operators cancel for the decay-term (as opposed to our numerical methods), we ignore these steps for the linear decay component to reduce bias (less spectral filtering). Our final model is thus:

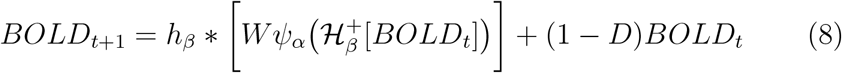

### 2.2. Task Model Derivation

Our approach leverages individualized resting-state models in order to estimate task-evoked brain effects, while making minimal modeling assumptions about the underlying task mechanisms. We model brain activity in task (*x_τ_*) as a dynamical system containing two components: an intrinsic dynamical component *f* (*x*) which is estimated from resting-state models (see previous section), and an exogenous input component *I_τ_*.

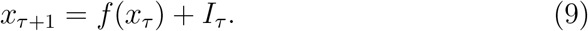

The latter component is exogenous with respect to the resting-state model and should not be interpreted as “exogenous to the brain”. Rather, *I_τ_* represents additional input to each brain region beyond that which is created through intrinsic (resting state) dynamics embedded in *f* (*x*). In principle, this technique is compatible with any resting-state model (*f* (*x_τ_*)). For the current work, we chose MINDy ([1], [17]) as it is highly scalable, nonlinear, and robust to many nuisance factors. The aim of the current work is to estimate the input (*I_τ_*) for task data and to investigate exogenous input as a marker for cognitive states. We do not assume a specific mechanism underlying this input (e.g. recurrent input, inter-regional signaling, neuronal “noise”, or sensory afferents are all possible sources) or any spatial/temporal properties of *I_τ_*. Thus, we treat *I_τ_* as a latent signal to be estimated (i.e., by filtering *I_τ_* from BOLD). By contrast, other methods, such as DCM ([19],[23]) assume a time course of *I_τ_* (the temporal aspects of *I_τ_*) based upon task design and only estimate its relative contribution to each brain area. For this reason, we term our objective MINDy-based Filtering. Although the mechanisms of interest (*I_τ_*) are modeled as neural, fMRI measures the hemodynamic BOLD contrast. For this reason, we use MINDy to simultaneously model neural dynamics and the hemodynamics which link neural events to fMRI measurements. We assume that BOLD signal recorded in task reflects the convolution (denoted “*”) of latent neural activity (*x_τ_*) with a region-specific Hemodynamic Response Function (HRF; denoted *h*) estimated from resting state data ([17]). Thus, for each brain region (parcel “i”) our model of task BOLD is:

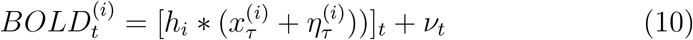

We consider noise at the level of the neurovascular coupling *η_t_* and at the level of BOLD measurements (*ν_t_*). These terms are modeled as normal random variables which are independently and identically distributed (iid) between brain regions and time points. Process noise (physiological stochasticity) is not explicitly modeled at the neural level in Eq. 9, as it is absorbed in the unknown inputs *I_τ_*. Substituting for *x_τ_* (from Eq. 9) and rearranging yields:

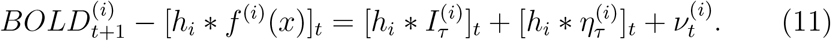

Thus, the HRF-convolved input [*h I*]*_t_* is equal to the difference between measured and predicted BOLD plus additional autocorrelated noise terms. For all current analyses we consider brain states estimated with HRF-convolved estimates of input ([*h I*]*_t_*) as opposed to the estimates of *I_τ_* alone. This step enables the same statistical pipelines (i.e. GLM structure) to analyze original fMRI BOLD data and the HRF-convolved input. As a result, the estimation of [*h I*]*_t_* serves as an additional “preprocessing” (filtering) step that can be added to any fMRI pipeline with minimal effort. No information regarding task events is used in estimating *I_τ_*, so the same statistical frameworks are applied to model-filtered and original data.

### 2.3. MINDy-based Filtering

In the current approach, we do not explicitly model different forms of noise. The only noise factor we consider is the measurement noise power in inverting BOLD onto neural activity. Since neurovasculature noise is removed (*η_t_*=0), Wiener deconvolution ([22]) generates the least-mean-square estimate for *x_t_*. The resultant approximation for BOLD-convolved input ([*h ∗ I*]*_t_*) is:

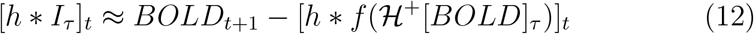

With 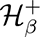[*BOLD*] denoting the Wiener deconvolution of each region’s BOLD signal with respect to the corresponding HRF model. Thus, we estimate neural activity by deconvolving BOLD with the region-specific HRF’s identified at rest. Predictions are made in terms of neural activity and then reconvolved to produce predictions in terms of BOLD. The difference between measured and predicted BOLD approximates the HRF-convolved input. All operations are performed over the whole timeseries simultaneously.

The full procedure is thus:

1. Resting-state data is used to estimate MINDy model parameters: connectivity (*W*), transfer-function curvature (*α*) and decay-rate (*D*) as well as the HRF shape (*β*). *ω* := *W, α, D, β* according to the dual model:

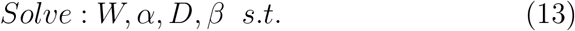

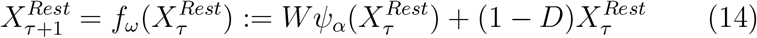

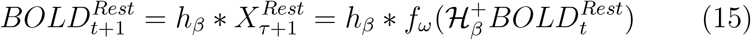
2. Using HRFs estimated from rest, measured BOLD-level task data is deconvolved to neural-level.

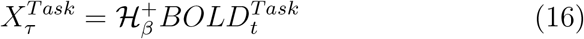
3. The parameterized MINDy models use deconvolved observations to predict task neural activity 1TR into the future.

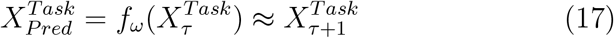
4. Predicted neural activity is convolved into predicted BOLD measurements.

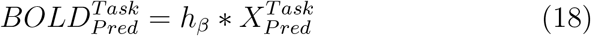
5. “Filtered” timeseries are calculated by subtracting the predicted future BOLD from measurements.

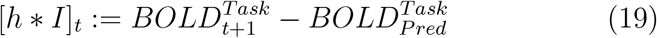

For the univariate-linear (decay) terms, analytic convolution and deconvolution cancel so we only performed these steps on the nonlinear terms to minimize bias (numerical implementations do not fully cancel). This choice also enabled direct comparison of brain-wide MINDy models with local auto-regressive models (see Sec. 3.10). Model predictions are thus:

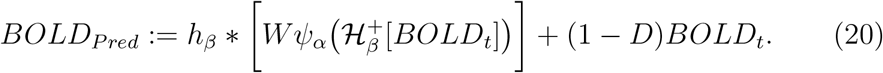

## 3. Methods

### 3.1. Subjects

Data consisted of fMRI task and resting-state scans for 71 healthy young-adult subjects collected as part of the Dual Mechanisms of Cognitive Control (DMCC) study ([24]). We note that the DMCC participant pool contains a large number of monozygotic and dizygotic twin pairs although this feature was not relevant for our analysis.

### 3.2. Scanning Protocol

Each participant took part in three separate scanning sessions which occurred on different days, but all had the same general procedure. Each day, participants provided two resting-state scans of 5 minutes each as well as two scans each for four cognitive tasks: the AX-Continuous Performance Task (AX-CPT), Sternberg Task, Stroop Task, and Cued Task-Switching (Cued-TS). The two scans per task were performed sequentially for each task whereas the two resting-state scans were separated in time (one at the session start and one at the session midpoint). Each of the task scans (2 per task per day) contained three task-blocks separated by inter-block intervals and lasted approximately 12 minutes. For resting state and task, the two scans per day were split between anterior-posterior and posterior-anterior phase-encoding directions. Scans were performed at 3T with 1.2s TR (multi-band *×*4; see [24, 25] for additional details).

### 3.3. Task Descriptions

We briefly describe the general structure of each of the four cognitive tasks in the “baseline” format which was administered on the first scanning day (see [24],[25] for more details on task design and rationale). Subtle changes to task structure were made on the two following days (subsequent section) but were not relevant to our analyses. The **AX-CPT** task ([26]) involves repeated sequences of cue-probe pairs, in which the response to the probe item is constrained by the preceding contextual cue. Thus, the A-X cue-probe pairing requires a target response and is frequent, leading to strong associations between the cue and probe. However, both the B-X pairing (where “B” refers to any non-X cue) and A-Y pairing (where “Y” refers to any non-X probe) require non target responses. In the **Sternberg task** ([27]), participants are sequentially presented with a short list of words to memorize for that trial (called the memory set; appearing across two encoding screens). After a short retention delay, they are presented with a probe word and must determine if the probe was present in that trial’s memory set. On some trials, the probe item is termed a “recent negative”, in that was not present in the current trial memory set but was present in the memory set from the preceding trial. In the current implementation of the **Stroop task**, subjects are asked to verbally report the font color in which probes are displayed ([28]). Each probe is itself a color-word, and can either be congruent (font color is the same as the color word, e.g., BLUE in blue font) or incongruent (font color is different from the color-word name; e.g., BLUE in red font). Lastly, during **Cued Task-Switching** (Cued-TS, [29]) participants are pre-cued to attend to either the number or letter component of a subsequent probe (combined letter + digit). In “attend-number” trials, participants indicate whether the digital component of a probe is even vs. odd. In “attend-letter” trials, participants indicate whether the letter component is a consonant vs. vowel. The probe can be either congruent (both letter and digit are associated with the same response) or incongruent (the letter and digit are associated with different responses). With the exception of the Stroop task, participants report responses using button presses.

### 3.4. Cognitive Control Demand

The current set of trial-based analyses center upon the ability to identify neural signatures of cognitive control. Although cognitive control is a heterogeneous construct, we specifically studied the conflict resolution aspects of cognitive control, so we use the terms control-demand and conflict interchangeably when referring to these tasks, and contrasts between trial types. In particular, we operationally identify cognitive control demand as the difference in neural activity measures during high and low-conflict trials for each task. In the AX-CPT, we contrast BX trials (high conflict) vs. BY (low conflict). The BX trials are high conflict because of the target-association with the X-probe, which require contextual cue information to override. For the Sternberg task, we contrast trials with recent negative probes (high conflict) and trials containing novel negative probes (low-conflict). Thus, recent negative trials are high conflict because the familiarity of the probe, requires information actively maintained in memory to override. In the Stroop task, we contrast incongruent (high conflict) and congruent (low conflict) trials. The incongruent trials are high conflict because the task goals (name the font color) are required to override the dominant tendency to read the color-name. Lastly, in the Cued-TS we also contrast incongruent (high conflict) and congruent (low conflict) trials. The incongruent trials are high conflict because it is critical to process the task cue, in order to know what response to make (for congruent trials, the same response would be made regardless of the task being performed).

### 3.5. Task Manipulations

The four tasks (AX-CPT, Sternberg, Stroop, and Cued-TS) were chosen to measure/engage cognitive control. On the first scanning day, participants performed a “baseline” version of each task. On the subsequent days, however, participants performed modified version of each task, meant to promote either proactive or reactive cognitive control strategies. On the two subsequent scans participants performed all the reactive-mode conditions of the tasks on one day and all the proactive-mode conditions of the tasks on another, with the order of proactive vs. reactive days counter-balanced across subjects. In the current work we do not consider the influence of cognitive-control mode and combine data for each task across scanning sessions, to increase statistical power.

### 3.6. Behavioral Measures

In each task we recorded two behavioral measures: reaction time (RT) and accuracy. Reaction times for button presses were recorded digitally, whereas reaction time for the Stroop task was defined by the duration of silence (time until participant begins a verbal response; see [24]). For the current work, we focused upon the difference in performance measures between trial-types with high cognitive control demand and those with low cognitive control demand (see below). As in previous work with these tasks, we observed lower performance (higher RTs and lower accuracy) on the high demand trials indicative of a cognitive control effect ([24]). For the RT data, we defined cognitive control effects as the difference in normalized RTs between high and low-control trials:

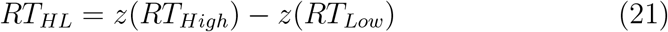

with *z* denoting z-score normalization. We separately normalized the high and low RT conditions to account for potential heterogeneity of variance between conditions. However, we could not separately normalize accuracy by condition as some of the low-control distributions were near-degenerate (e.g. in one Stroop session over 90% of subjects had 100% accuracy for low-control trials). Similarly, we obtained near-identical results using either the high vs. low contrast for accuracy and using just high-control trials (since low-control accuracy was near-ceiling). For parsimony, we chose to use the high-control data for plots as opposed to the near-identical high vs. low contrast. As with neural data, we averaged the normalized response times between sessions for each task. Interestingly we found that, unlike RTs, neural data using conventional techniques only predicted accuracy in the baseline session. Therefore, we only used the baseline accuracy for benchmarking (averaged over tasks) and similarly for neural data.

### 3.7. Pre-processing and Parcellation

Raw resting-state and task data were preprocessed using the same pipeline, implemented with fMRI-prep software ([30],[31]). The whole-brain surface data were then parcellated into 400 cortical parcels defined by the 400 parcel Schaefer atlas (Schaefer [32]; 7-network version). Subcortical volumetric data was divided into 19 regions derived from FreeSurfer ([33]). Motion time-series consisted of the 3-dimensional coordinate changes for rigid-body (brain) rotation and translation (6 total). Motion and linear drift were regressed out of pre-processed resting-state data before MINDy model fitting and from task data prior to filtering. Since motion time-series are also covariates within our task GLMs (as is common), this step does not bias results, as motion is implicitly removed from the unmodeled data during GLM estimation (see below). However, we also implemented controls (see Sec. 3.10) which used this same data (i.e. motion pre-regressed) with conventional analyses.

### 3.8. Task GLM Analyses

Statistical models of task fMRI were estimated using general linear models (GLM) as implemented in AFNI. The same analyses were performed for all data pipelines (e.g. original and MINDy-Filtered). Two classes ofGLM were used for each task: one designed to estimate event-triggered effects and another to estimate sustained activity. These models only differ in that the event-related GLM models contain separate terms (FIR models) for each trial-type whereas the sustained GLM does not distinguish between trial-types, which enabled better estimation of the sustained effects (block regressor). The GLM design consisted of a mixed block/event-related design in which trial-type effects were modeled using a modified Finite-Impulse-Response (FIR,[34], [35], [36]) framework (AFNI TENT; [37]), whereas block effects (task vs. inter-block interval) were modeled using a canonical HRF convolved with the block regressors. The TENT bases (“knots” in AFNI terminology) generated an FIR design with each basis representing one TR (relative task start). The GLM design also included block onset/offset (modeled with a canonical HRF) and the six motion regressors corresponding to rigid body translation and rotation (3 each). Timepoints containing excessive motion (Framewise Displacement *>* 0.9mm) were censored from task GLMs. Estimation was performed using the built-in AFNI function “3dREMLfit”.

### 3.9. MINDy Modeling

Mesoscale Individualized NeuroDynamic (MINDy, [1][17]) models were generated from each subject using the parcellated, pre-processed resting-state data for each subject, combined across scanning sessions. Thus, a single MINDy model was estimated for each subject and was used in analyzing task-data across scanning sessions. We simultaneously estimated the neurovascular coupling/HRF and latent brain networks by combining the original MINDy model with Surrogate Deconvolution as in [17]. This combination simultaneously estimates HRF kernel parameters for each brain region as well as the connectivity matrix, region-specific transfer function shape, and local decay parameter (time-constant). Previous work indicates that the inclusion of Surrogate Deconvolution renders MINDy estimates robust to spatial variation in the HRF. Moreover, the spatial distribution of estimated HRF properties such as time-to-peak are consistent with empirical literature at the group level and are also reliable at the level of individual differences ([17]). Hyperparameters used in MINDy model fitting were identical to previous studies ([1]), but with batch sizes decreased to 150 TRs each in order to accommodate the shorter scan lengths of this dataset.

### 3.10. Control Pipelines

In addition to comparing the proposed pipeline with conventional analyses, we also repeated all task analyses for several control pipelines (Fig. 1B). These control pipelines considered two factors that might explain results: 1) pre-processing and 2) mechanistic components of the model (SI Sec. 7.7). The MINDy modeling framework assumes that nuisance covariates such as motion and drift have already been removed from time-series prior to model fitting. Therefore, to address #1, we implemented a control in which standard GLM analyses were computed directly upon the fMRI BOLD task timeseries, with motion covariates already regressed out first. The same regressors also appear in the task GLM model (which is shared across all pipelines), but regressing these factors out first will rescale estimated beta-coefficients due to the input normalization performed by many fMRI processing packages (e.g. AFNI). This control ensured that improvements in group-level sensitivity were due to increased similarity of estimated spatiotemporal patterns rather than theoretically uninteresting factors due to pre-processing pipelines. We refer to this control as “pre-regressed” (pre-Reg). Estimates using this pipeline were nearly identical to the original pipeline and event-related coefficients were highly correlated (average over tasks: *r* = .97), collapsing over subject, parcel, and TR during the probe period.

In the SI (Sec. 7.7), we address #2 by considering the influence of anatomically local dynamics vs. interactions between brain regions. This distinction is significant for three reasons. First, it is theoretically significant to distinguish between purely local neural dynamics and inter-regional brain dynamics. Secondly, long distance interactions between brain regions cannot be explained solely in terms of neurovasculature since the regions involved may share anatomically distinct blood supply (i.e. different cerebral arteries). As a result, improvements identified in whole-brain models, but not purely local models, cannot be explained solely as a benefit of hemodynamic modeling (although other contaminants such as motion could still be a factor). Lastly, analyses using the purely local models are equivalent to region-specific frequency-domain filtering. Although this equivalence does not imply that neural dynamics are insignificant, the signal-processing interpretation is simpler and could render the proposed neural modeling framework unnecessary (i.e. less parsimonious). Thus, the local dynamics control serves to ensure that our guiding neural modeling framework provides additional value above its (partial) relationship to existing signal-processing techniques. This control was implemented in two distinct variants: either heterogeneous (region-specific) or homogeneous (region-invariant) autoregressive models fit to each subject.

The homogeneous model consists of an autoregressive model that is specific to subject, but not parcel:

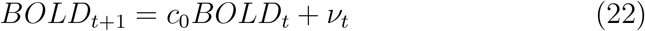

We assumed that the noise-component was independent and identically distributed between parcels and solved for *c*_0_ using linear regression (collapsing BOLD across parcels). The “input” estimates from this model consist of the residuals (*ν_t_*). We fit the heterogenous model analogously to the homogeneous model, but with region-specific autoregressive terms:

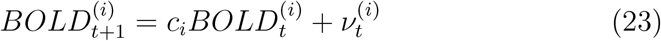

for parcel “i”. We use these two cases to determine whether regional heterogeneity is a significant factor in any improvements due to local modeling. We refer to the homogeneous and heterogeneous models as global (“glob”) and local (“loc”) autoregressive (AR) models, respectively. Results were generally similar for the two AR models (high-low coefficients correlated *r* = .99)

## 4. Validation and Comparison Criteria

In order to assess potential advantages of MINDy-based Filtering, we considered two types of comparisons: benchmarking (is method “a” better than “b”?), and sensitivity/robustness (how does factor “x” influence method “a” vs. “b”?). The first case establishes whether MINDy-based Filtering offers additional statistical power in detecting task effects. The second case establishes whether MINDy-Based Filtering enhances statistical power for detecting task effects in a selective (i.e., to the regions showing significant task effects to begin with) or more global manner.

### 4.1. Benchmarking Event-Related Effects

Trial-types were defined by high cognitive control demand vs. low cognitive control demand across the four tasks (see Sec. 3.4). Trial-specific activity was modeled using a Finite Impulse Response (FIR) model with 1TR resolution (1.2s) and task-specific length (see Sec. 3.8). Group-level statistics were compared for the peak effect (parcel × method specific) over a task-specific 2TR interval. This interval was chosen during study piloting using the peak times in conventional analyses (starting from 1: AX-CPT:TR 7-8, Cued-TS: TR 8-9, Stern: TR 11-12, Stroop: TR 3-4). Thus, the analysis targets are statistically biased *against* the proposed technique since they were chosen to maximize conventional analyses. These times qualitatively correspond with a typical HRF time-to-peak after the probe-events which define high vs. low control trials (see Sec. 3.4). Previous literature and present results suggest that these effects are primarily one-sided, with activity increased in the high-conflict (control demand) trials relative to low-conflict (low control demand) in relevant brain regions (e.g. Fig. 2A). Conversely, task-negative effects (significant decreases) have largely been associated with sustained signals as opposed to high vs. low control events. For these reasons, we only considered significant increases in activity for trial-type analyses. Group-level t-tests (within parcel) were compared for all parcels with significant increases (either method; Fig. 2B), or for a set of 34 parcels (pre-defined from independent conventional analyses which showed consistent control-demand effects across all tasks, (Fig. 2A, SI Table 1, [24]). Since these parcels were pre-selected based upon conventional analyses, they are statistically biased *against* the proposed method (i.e. in favor of conventional methods).

**Fig 2.**
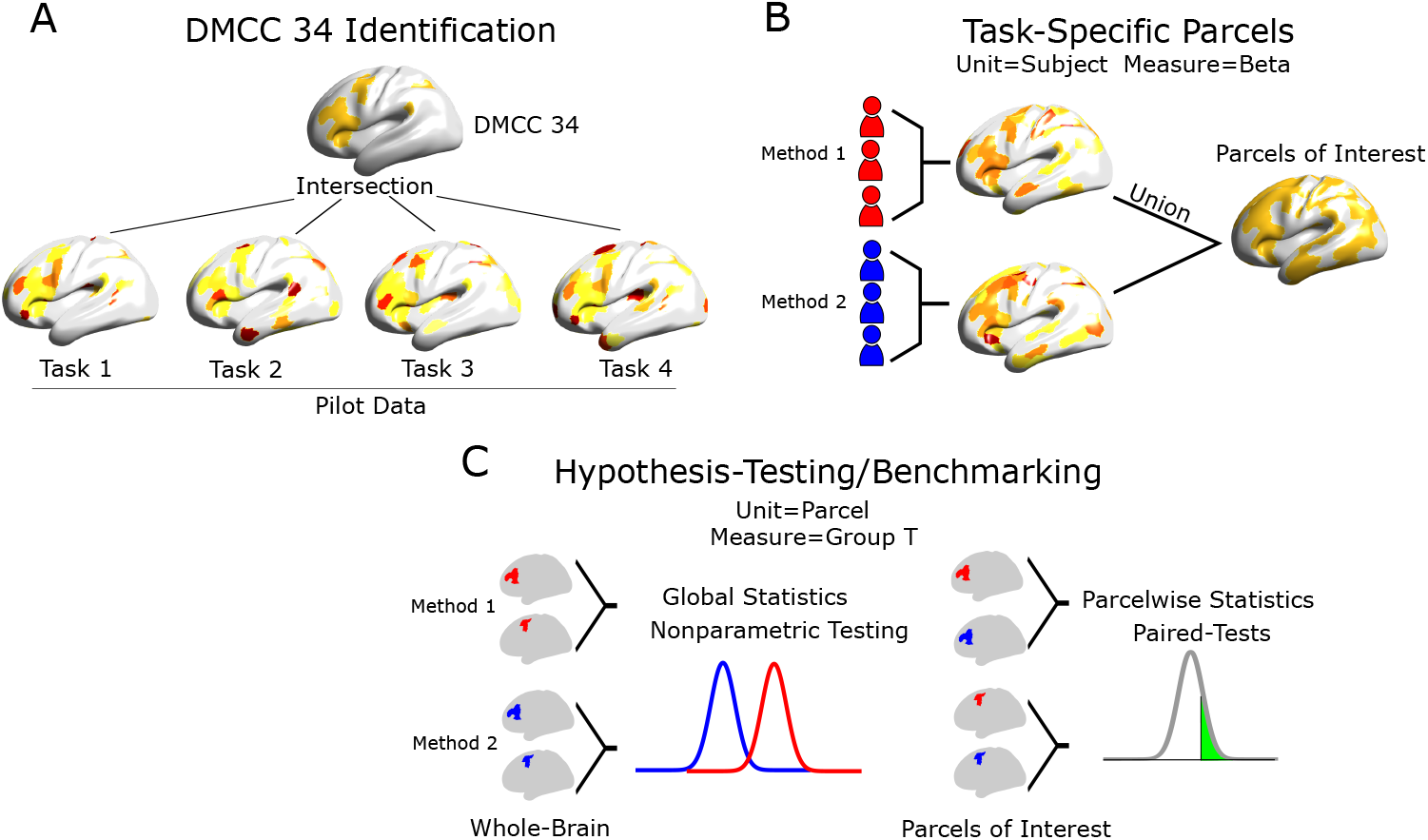
Statistical frameworks for comparing methods. A) The DMCC34 set of parcels was defined by all parcels which displayed an effect of cognitive-coontrol demand in every task based upon separate pilot data using conventional analyses. Hence, the DMCC34 set of parcels is pre-specified used across all tasks. B) Candidate regions for task-specific comparisons (parcels-of-interest) are identified for each pairing of task pipeline by combining parcels with group-T meeting *p < .*001 for at least one pipeline in a comparison (one-tailed for events, two-tailed for sustained effects). C) Data is analyzed either using resampling statistics for global measures (e.g. for brain-behavior correlations, generalizability) or in terms of paired-differences between methods over each parcel-of-interest.

**Table 1.**
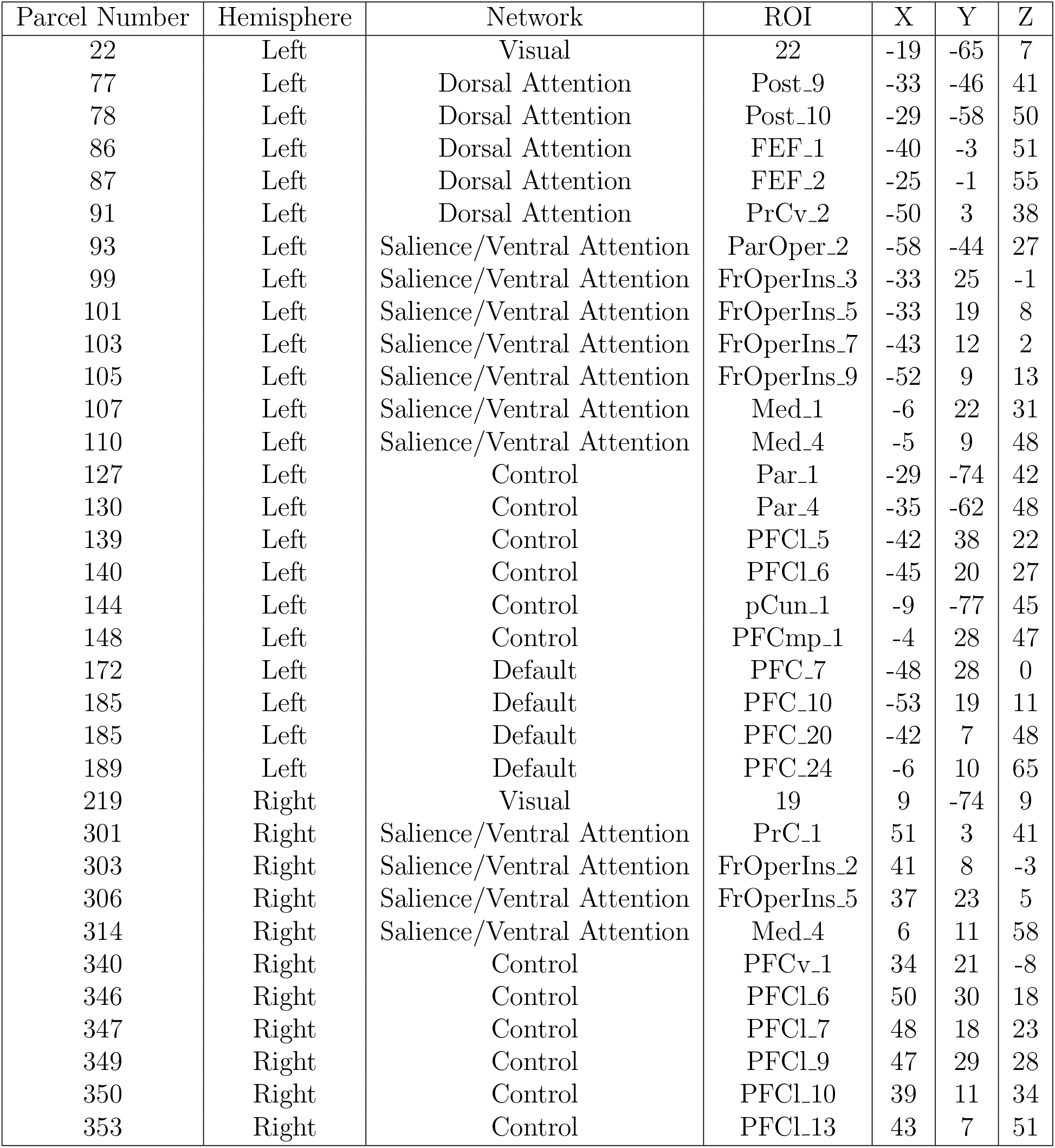
Attributes of the DMCC34 parcels. Indices are for the Schaefer 400-region, 7-network parcellation [32]). Coordinates (X,Y,Z) refer to MNI centroids.

### 4.2. Benchmarking Sustained Effects

In addition to event-related analyses, we also considered the identification of sustained effects (block-related changes). Results of these analyses are primarily presented in the SI (Sec. 7.5). As with event-related analyses. Sustained effects in a mixed block/event design refer to “background” activity that is present regardless of whether participants are performing a task ([38], [39]). Since we used FIR models to span each trial type, sustained effects in our analysis *only* refer to activity during inter-trial periods (non-trial periods of task-blocks) since effects during other periods are absorbed in the trial FIR vs. rest-block contrasts ([38], [39]). We compared the group-level effect size of each technique (MINDy-based Filtering and several controls) in detecting sustained effects. Methods were compared pairwise, and benchmarking analyses were only conducted on parcels which had a significant effect for either method in a pair. Sustained analyses considered both signal increases and decreases, so the target metric was absolute t-value (1-sample group test) for the GLM sustained betas (see Sec. 3.8).

### 4.3. Testing Selective vs. Global Improvements

We further analyzed benchmarking results by testing how MINDy-based Filtering changes the distribution across parcels. The primary question was whether the MINDy-based Filtering: a) uniformly changes statistical power across the brain (by shift or scale); b) primarily identifies previously insignificant regions or c) primarily alters previously significant regions. This analysis is important for determining whether the technique globally improves statistical power or differentiates task-relevant regions from the rest of the brain. We test these effects using multilevel linear models to compare MINDy-based Filtering to the different control models. These multilevel models (presented in more detail later) contain task-specific main effects of method (anatomically global) and additional terms for task-implicated (statistically significant parcels). We use these models to test the significance of model improvements (increased effect sizes) after discounting anatomically global changes.

### 4.4. Sensitivity to Cognitive States

Sensitivity analyses were performed to assess the impacts of cognitive states, individual differences, and motion. In the current case, cognitive states differ between tasks and trials. Although, each of the four tasks are commonly used to index cognitive control, cognitive tasks are not construct-pure. For instance, tasks featuring delays (AX-CPT, Cued Task Switching, and Sternberg) are thought to be more dependent upon working memory than those without delays (i.e. the Stroop task). However, many task-specific factors are the same between high and low control trials of the same task (i.e. all events prior to the probe). Thus, we control for cognitive similarity across tasks by comparing results across increasing levels of cognitive similarity: low-control trials, high-control trials, and the contrast high vs. low control trials. These levels increasingly isolate the cognitive control construct by increasing control demand (high-control trials) and controlling for other task events (high vs. low contrast). Methods which are sensitive to cognitive states will produce more similar results between task contexts when the cognitive states measured are more similar. Put simply, we studied between-task similarity in the whole-brain activation profile under the premise that more similar task conditions should lead to more similar activation.

We quantified similarity in the activation profile using the Intraclass Correlation (ICC; [40]) which generalizes the concept correlation to multiple groups (i.e., four tasks as opposed to pairs). Tasks differed in effect magnitude and there was no theoretical basis for assuming this factor should be identical between tasks (i.e. we don’t assume each task equally taxes cognitive control), so we normalized the group-average data (divided by the standard-deviation over parcels) for each task × method before using ICC to test similarity in activation.

### 4.5. Significance Testing for Construct Identification

We used permutation statistics to compare the significance of generalizability tests between methods. When testing the generalizability of group-level patterns, we treated brain regions as the object of measurement in intraclass correlations (ICC, [40]) over task classes. Larger ICC values imply more similar whole-brain activation profiles between tasks. We estimated confidence intervals with bootstrap sampling over the set of brain parcels.

### 4.6. Robustness to Motion

In an SI analysis (Sec. 7.8), we compared methods in their robustness to motion confound. While previous work has established that the model-fitting technique (MINDy) is robust to motion ([1]) it remains unknown whether MINDy-based Filtering technique also exhibits similar motion robustness. Therefore, we compared methods in terms of sensitivity to motion artifact. We considered three motion metrics for task data including the number of frames censored based upon framewise-displacement (FD) criteria (*<* 0.9mm), the median framewise displacement, median-absolute-deviation (MAD) of DVARS ([41]). We analyzed sensitivity by comparing the similarity (ICC) of results between high-motion and low-motion groups of subjects (median split for each motion measure).

## 5. Results

### 5.1. Structure and Presentation of Results

We designed analyses to answer four questions: 1) do resting-state MINDy models (partially) generalize to task? 2) does the proposed technique improve power in answering cognitive-neuroscience questions? 3) can these methods test hypotheses which were previously impractical? And 4) do improvements reflect theoretically interesting concepts (e.g. signal propagation) or do they stem from signal-processing/filtering side-effects? The first question resolves whether the intrinsic dynamics modeled at rest meaningfully generalizes to task (although not perfectly, as we are interested in the task versus rest differences). The second and third questions identify methodological contributions, whereas the last question addresses whether these techniques also offer additional theoretical insight (i.e. their success reflects some principle of brain function). This question is important for determining whether results reflect brain network dynamics or can be more parsimoniously explained in terms of (non-neural) signal processing effects.

In the main text, we emphasize comparing methods in event-related analyses due to the popularity of event-related designs. However, we also compared methods for the analysis of sustained-effects in a mixed block/event design. These results are presented in SI Sec. 7.5 and 7.6. We also tested the specific contribution of modeling connectivity by comparing MINDy-based Filtering with analogous filters using reduced (autoregressive) models (SI Sec.7.7).

### 5.2. Identification of Task-Relevant Parcels

In order to compare methodologies (“third-level” analysis) we first identified task-relevant parcels over which to guage improvements. We performed this step in two ways: either using a set of parcels consistently engaged across tasks (“DMCC34”) or separately identifying relevant parcels for each analysis (i.e., for the different tasks; Fig. 2A,B). In the first case, we used pilot data and conventional analyses to identify a set of 34 brain regions which displayed significant increases (*p < .*05, Bonferoni-corrected) in activity due to cognitive-control demand across all four tasks (Fig. 2A). This set is referred to as “DMCC34” and constitutes a “pre-specified” comparison set as it was developed using a separate set of pilot subjects. It is also biased away from finding MINDy-based Filtering improvements, since, by definition, the parcels were identified as maximizing conventional univariate statistical contrasts.

In addition, we identified “parcels-of-interest” specific to each third-level comparison (i.e., task + methods; Fig. 2B). We defined “parcels-of-interest” as reaching an uncorrected significance of *p < .*001 for at least one of the methods being compared (Fig. 2B). We used a slightly more liberal criteria for identifying these parcels as several of our “third-level” analyses compare second-level analyses over parcels-of-interest (Fig. 2C), although we later demonstrate that general improvements in detection power hold across significance thresholds (Sec. 5.5). These “parcels-of-interest” are also specific to a given second-level contrast (separate sets for events and for sustained (block-related) effects). Thus, for each pair of methods (e.g. MINDy vs. original) we identified one sustained and one event-related set of parcels for each of the four tasks.

### 5.3. Resting-state Model Predictions Generalize to Task

The key premise of our approach is that task effects are marked by systematic deviation from intrinsic brain dynamics, reflecting extrinsic influences (“input”). As such we seek to estimate these influences by filtering out intrinsic dynamics to recover task “input” (we stress that “input” should not be taken literally; see Sec. 2.2,6.3.2). In practice, this operation corresponds to computing the difference between model-predicted and observed changes in brain activity at each timestep. The validity of our framework thus rests upon three claims: 1) that task events are marked by (slight) deviations from intrinsic-dynamics, 2) that these deviations are systematic and can be modeled as additive “input” to the otherwise preserved dynamics, and 3) estimated inputs are a more consistent marker of task effects than the original BOLD signal.

Our first claim, that task events deviate (slightly) from intrinsic dynamics is observed by comparing MINDy prediction accuracy over task and “rest” blocks (3 task blocks and four rest blocks per run). During “rest” periods, prediction accuracy is nearly as high as for the training resting-state data. Overall, the range of model prediction accuracies for resting-state scans (*R*^2^ = .58 ± 06) was roughly similar to that observed during task (*R*^2^ = .56 ± .08*, .*54 ± .07*, .*56± .08*, .*50 ± .09, for AX-CPT, Cued-TS, Stern, and Stroop, respectively; Fig. 3A). However, prediction accuracy differed between periods in-between task blocks (“rest” blocks) and when subjects were actively engaged in task. During “rest” blocks, MINDy predictions were no worse than for resting-state scans. In AX-CPT and Stroop accuracy during “rest” blocks was significantly greater than for resting-state scans (*paired* – *t*(70) = 3.5*, p* = .0008; *paired* – *t*(70) = –4.5*, p* = 2.4*E* – 5) and for the other two tasks, the MINDy modeling of resting-state scans and rest-blocks within task scans was equally accurate (*t*(70) = –1.1*, t*(70) = 1.2*, n.s.*) for Cued-TS and Sternberg). By contrast, model accuracy decreased when subjects were actively performing each task (*p^/^s* ≤*E* − 8), while remaining well above chance (*R*^2^ = .54± .08*, .*52± .08*, .*54± .08*, .*45± .10, same task order; Fig. 3A,B). An illustration of the pattern is shown for a representative task (Cued-TS), showing the amount of variance (*R*^2^) explained by MINDy at each TR across the whole-scan timeseries (Fig. 3B). Deviations from model predictions (unexplained variance) are also greatest during the probe/response period (Fig. 3C), indicating that these deviations are a strong marker of task events. Thus, intrinsic dynamics observed at rest still predict task dynamics, but the degree of accuracy is tightly coupled to task events.

**Fig 3.**
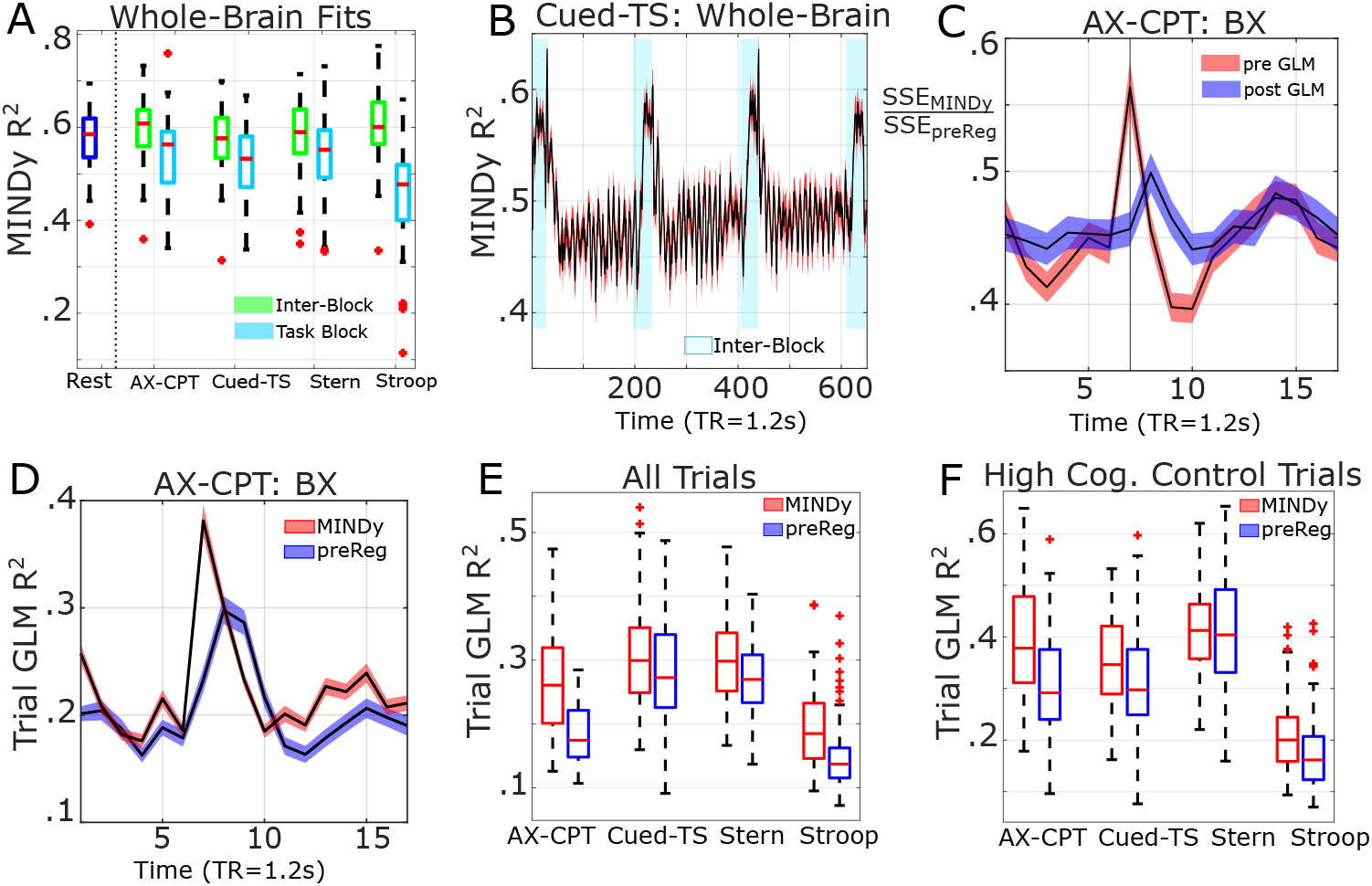
Validation of MINDy-based Filtering Framework. Task effects are defined by deviation from intrinsic dynamics. A) Intrinsic dynamics modeled by MINDy with resting-state data, remain valid (but less accurate, see SEC) in task. B) Deviation from intrinsic dynamics (i.e., estimated “input”) mark periods of active task engagement over long timescales (task blocks) and C) short-timescales (task events; pre-GLM). There is a peak in unexplained event-related variance (SSE MINDy; pre-GLM) timed to the onset of probe effects. However, this variance is well-explained by task GLMs (post-GLM) indicating that event-related deviations from MINDy (fit to rest) are well-described as additive “input” to the model. D-F) Timeseries post-MINDy based filtering (red) have a greater proportion variance attributed to task events. Statistics are averaged over a set of pre-specified parcels-of-interest (DMCC34). D) Example timecourse from a BX (high-control) trial in AX-CPT demonstrates clear increases in task-explained variance during the probe-response period (7-8 TR). E) MINDy-based Filtering significantly increased signal variance attributed to any task event in four tasks. F) Improvements in high-control trials were significant in 3 of 4 tasks (all but Sternberg). Shading indicates standard error over subjects. “post-GLM” indicates that both the numerator and denominator SSE are taken after performing GLM (MINDy=MINDy-Filtered), whereas “pre-GLM” indicates the relative sum-of-squares after MINDy-based Filtering but before fitting task GLM models. Both B) and C) are taken from AX-CPT (averaged over scans). Time-courses in C) are event-locked to the start of “high-control” trials. Vertical line indicates TR7 which marks probe-related effects in AX-CPT (TR 7-8). “MINDy” denotes results using MINDy-based Filtering, while “pre-Reg” denotes the pre-regressed control (conventional analyses, but with additional motion-regression performed pre-GLM).

Our second claim is that these deviations are systematic and can be well-approximated as an exogeneous “input” to the existing dynamics. Statistically, this assumption corresponds to the residuals (MINDy prediction minus observed) being shifted (event-locked change in mean) during task events, as opposed to changing variance, which could reflect a breakdown of the underlying dynamics. For this analysis we only considered parcels known to be task-related: the DMCC34 set as the subsequent analyses assumes that the signal is task-related. Using Finite-Impulse-Response GLM designs we compared residual sum-of-squares before and after removing the effect of trial-period. Squared errors were averaged over the DMCC34 parcel-set for each subject. Analyses demonstrate that the probe-related increase in error (task-average: *t*(70) = 7.2*, p <* 4*E* − 9) is captured by an additive main effect of trial-period as the post-GLM unexplained sum-of-squares was not greater for the probe period in any task (n.s. 1-tailed) and actually decreased overall (task average: *t*(70) = −4.5*, p* = 2.6*E* − 5). Thus, task-induced deviations from intrinsic dynamics are systematic and well-described by additive “input” to the system.

Lastly, we assume that removing (“filtering”) intrinsic dynamics will accentuate task effects in the data by removing variance due to intrinsic dynamics. At present, we only consider spatially univariate effects (unlike e.g., MVPA), hence we tested the relative variance explained by task with and without MINDy-based Filtering. As in the previous analysis, we used the mean over DMCC34 parcels, as this analysis assumes that there is a true task effect to accentuate. Results indicate that MINDy-based Filtering generally increased the variance associated with task events. This result held for all tasks when combining across trial-types and for three-of-four tasks (all but Sternberg) when restricted to high-control trials. Thus, MINDy-based Filtering has the potential to improve the variance associated with task effects in human BOLD. We note that some inter-trial variability in brain activity can be related to behavior, so future study is needed to understand how MINDy-based Filtering affects veridical trial-to-trial variation (in a later section we find improvements in inter-subject behavioral prediction). However, these results demonstrate that our approach is well-justified and statistically powerful in identifying the types of simple (univariate) models of brain activity that are most common in neuroimaging.

### 5.4. MINDy-based Filtering Accounts for intra and inter-subject Variability

We also tested whether these intrinsic dynamics explain unique variability above the task GLM. This test is important for determining whether MINDy serves to predict the mean brain-response for each trial-type or whether it also predicts trial-to-trial variability. We quantified these properties through sum-of-squares partitioning (ANOVA). Across all tasks, we found that the proportion of unique variance explained by MINDy was significant (41.2% on average, Fig. 4A). However, MINDy predictions and the task effects do have some overlap (a non-zero MINDy×task sum-of-squares, Fig. 4A), thus MINDy predictions account for some of the variation in both the trial-to-trial variability (variation unique to MINDy) and the typical response across trials (MINDy *×* task interaction). We also tested how MINDy-based Filtering impacts variability in the evoked-response between subjects. We restricted these analyses to the pre-defined set of regions (the DMCC34 parcels, [24]) which were previously identified as having a significant control-demand effect across tasks. Results demonstrated that MINDy filtering decreased inter-subject variability in both main effects of trial-type (e.g. Fig. 4D) and the contrast between trial-types (e.g. Fig. 4E). In particular, these analyses and associated event-related timecourse visualizations reveal that the peak task-related effects become sharper (more well-defined), as well as more temporally-precise, after MINDy-based filtering. We used ANOVA to partition variance in the cognitive control effect into group-level variance and individual variance over the relevent (probe) trial periods.

**Fig 4.**
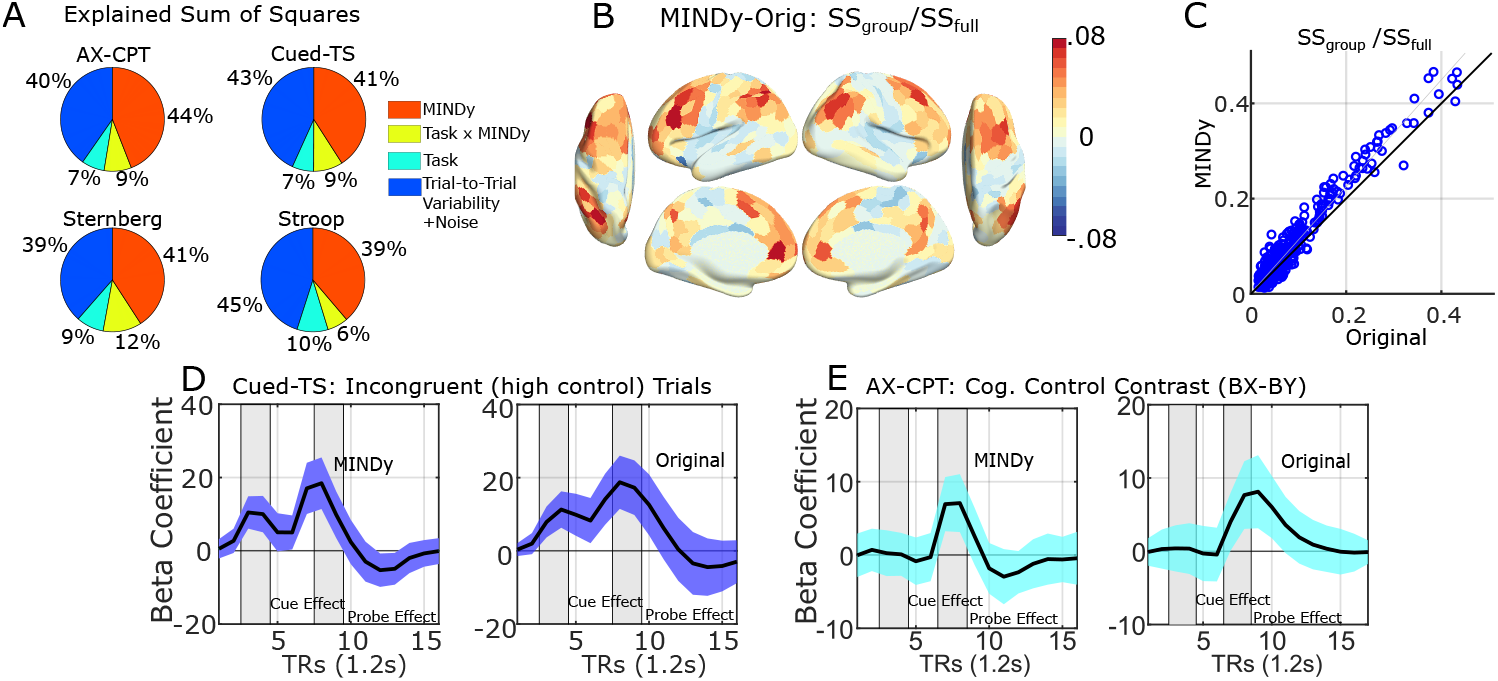
MINDy-based Filtering reduces variability within and between subjects. A) MINDy-based Filtering accounts for a significant portion of unique variability within each subject’s data. This effect holds across tasks (results averaged over all parcels, subjects). Variance partitioning was performed after removing variation due to nuisance factors (motion and drift). B) Difference in the relative group-explained variability between MINDy and the original data. Note that MINDy-based filtering actually decreases the proportion of group variance in some regions, but increases for task-implicated regions (e.g. lPFC). C) Group-explained variability particularly increased in parcels which already had a strong effect under original analyses (putative task-relevant parcels). D) MINDy-based Filtering reduces the between-subject variability of task-evoked signals. Example shown is the mean signal over the DMCC34 parcels for the Cued-TS high control-demand condition (incongruent trials). E) Variability also decreases for contrasts between conditions. Example shown is for the AX-CPT (BX-BY contrast). ‘MINDy” denotes results using MINDy-based Filtering before performing GLM, while “Orig” denotes the conventional pipeline (no MINDy).

We then tested whether MINDy increased the proportion of cognitive control effects attributed to a common group factor (sum-of-squares explained for the group model divided by the full/subject-specific models). As expected, regions implicated in cognitive control, such as the lateral and medial prefrontal cortex, anterior insulae, posterior cingulate, and posterior parietal cortex, had larger proportions of variability explained by the common group factor (analogous to Fig. 5A,B). MINDy-based Filtering increased the proportion variance explained by group-level models (relative full models) for the DMCC34 parcels (Δ*µ* = .034 ± .023*, paired − t*(33) = 8.56*, p* = 6.9*E −* 10). Brain-wide, parcels in which MINDy increased group-explained variance, had larger group-explained variance in the original analysis (*t*(417) = 4.92*, p* = 1.2*E* − 6) and the increase in group variance-explained (MINDy-Orig) was correlated with the original variance explained (*r*(417) = .40*, p* = 7.5*E* − 17). Thus, MINDy-based Filtering only increases group-level effects in task-implicated brain regions (those that already had a group-effect). Conversely, the relative variance attributed to subject decreased correspondingly (same statistics, but sign-flipped since *SS_Indiv\Group_/SS_Full_* = 1 − *SS_Group_/SS_Full_*). Thus, by removing intrinsic brain dynamics, MINDy-based Filtering reveals more similar task-effects between subjects.

**Fig 5.**
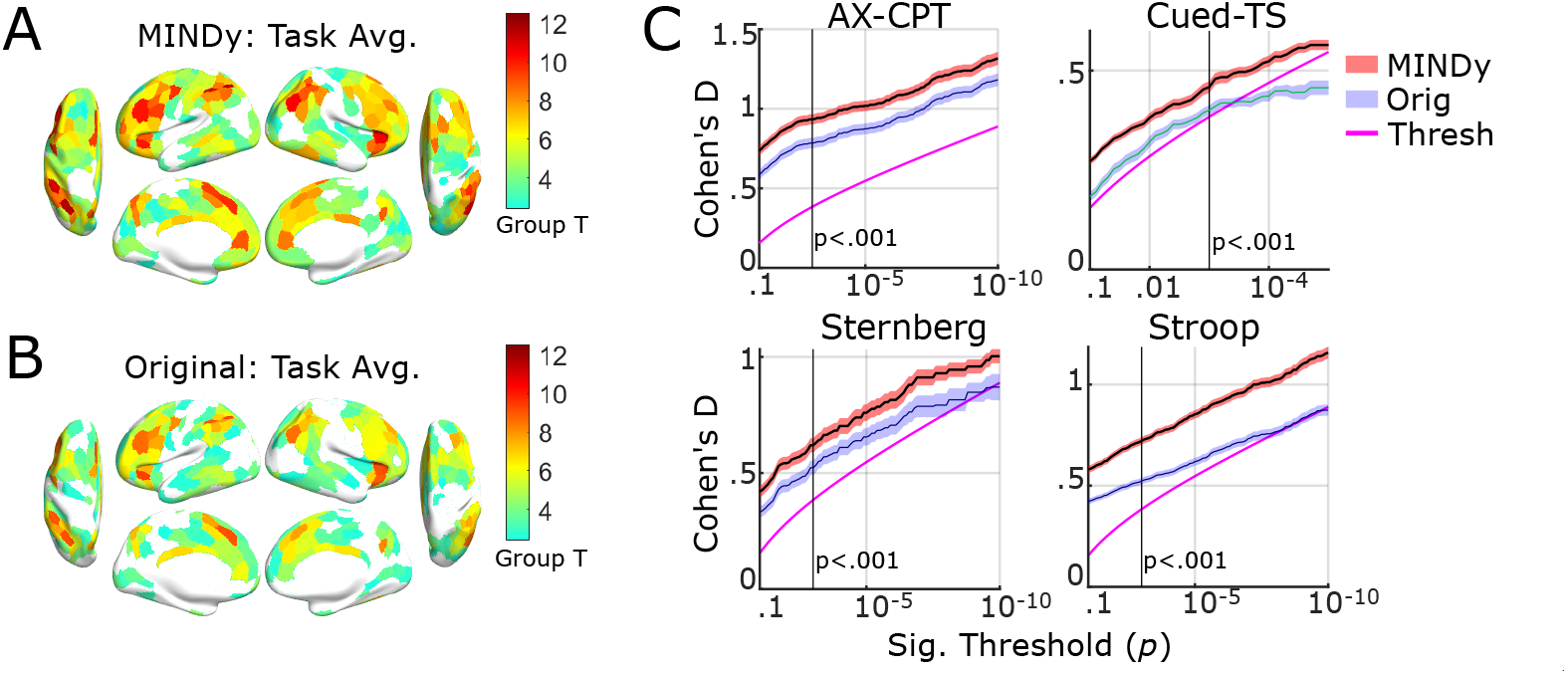
MINDy-based Filtering improves statistical power in identifying task effects. A) Average group-level *T* -statistic for MINDy-based Filtering across tasks in which the parcel had a significant cognitive control effect for at least one method. Uncolored parcels did not meet significance averaged across tasks. B) Analogous results for conventional analyses. C) Effect-size (Cohen’s D) for parcels meeting significance for at least one method by task and across significance thresholds (uncorrected). Magenta indicates the corresponding thresholds in terms of effect size (one-tailed) and shading indicates standard errors. “MINDy” denotes MINDy-based Filtering and “Orig” denotes the original pipeline. We only plotted the original pipeline for comparison due to visual overlap with results from the pre-regressed pipeline (i.e. original and pre-regressed were indistinguishable).

### 5.5. Improved Group-Level Detection Power

We tested whether MINDy-based Filtering improved statistical power in detecting group-level neural effects for each task, and in an omnibus test across tasks (Fig. 5A,B). For each event-related pairwise comparison of methods, we calculated group-level statistics from the GLM beta estimates of each task-relevant parcel (see Sec. 5.2). Results indicate that MINDy-based Filtering significantly increased statistical detection power on all tasks (four of four) for the event-related contrast relative to both the traditional pipeline and the pre-regressed control pipeline (all *p*’s≤1.2E-4; Fig. 5 C). For omnibus analyses, we collapsed observations across tasks (Fig. 5A,B). Results indicated that MINDy-based Filtering generally increases statistical power for event-related analyses (vs. original: paired-*t*(495) = 27.5, *p ≈* 0, vs. pre-regressed: *t*(492) = 27.9, *p ≈* 0).

We also tested whether improvements depended upon the criteria used to select task-relevant parcels, since methods were only compared on these parcels. Whereas the previous analysis used a fixed selection criteria (see Sec. 5.2), this analysis compared methods over a range of statistical thresholds for identifying task-relevant parcels to ensure results generalize across dietection criteria. Thresholds were defined by uncorrected within-method (second-level) significances ranging from *p* = .1 to *p* = *E -*10, one-tailed. We compared methods on all parcels that met a given threshold for at least one pipeline (original, pre-regressed, or MINDy). We imposed a minimum of 5 parcels for comparison which restricted the range of Cued Task Switching (minimum threshold: *p* = *E* − 5), while all other tasks had a sufficient number of parcels (AXCPT: n=10, Stern: n=7, Stroop: n=58) meeting even the most stringent criteria (*p* ≤*E* − 10). Results indicated that MINDy-based Filtering improved statistical power (effect size) relative conventional analyses on all tasks for all detection levels considered. Our approach also increased statistical power relative the pre-regressed control for all but one case (when only five parcels were compared for Cued-TS; *t*(4) = 2.5*, p* = .065, 2–*tailed*). We conclude that the proposed techniques improves statistical power in task-related parcels, regardless of how strictly “task-related” is defined.

One limitation of the previous tests, however, concerns the determination of which parcels are included in analysis: we compared effect sizes in parcels that met a significance criteria (i.e., already had large effect sizes). This approach is anatomically parsimonious in that the comparison regions are informed by data rather than prior assumptions. However, this dependency could produce biases. Therefore, we repeated the previous analyses over a fixed set of 34 pre-specified brain parcels (SI Table 1, [24]) that demonstrated significant increases due to cognitive conflict (event-related contrast) across all four tasks during independent and pre-specified analyses (see Methods, [24]). The implicated parcels agree with previous studies mapping the neuroanatomy of cognitive control and are largely located along lateral prefrontal cortex and anterior insula (Salience/Ventral Attention and Control networks; [42], [32]). Analyses over this restricted, pre-specified group of parcels agreed with the previous results: the omnibus (all task) statistical detection power and the task-specific effect sizes all improved relative to both the original pipeline and the pre-regressed controls (maximum *p* = 1.8*E* − 4). Thus, results indicated that MINDy-based Filtering improved statistical detection even when analyses were restricted to this group of 34 pre-specified parcels.

### 5.6. MINDy-based Filtering Selectively Enhances Task-Related Neural Signals

Results in the previous section indicate that the proposed technique increases the statistical detection power of task effects (Fig. 5C). Statistical power and effect sizes are useful benchmarking criteria as they are easy to interpret and relate to potential applications. However, these markers are also limited in that they indicate the ability to reject a generic null hypothesis of no task effects. Yet this generic null is not always a useful benchmark from which to provide additional scientific insight. For instance, approaches which magnify anatomically global effects may provide little benefit to functional “brain-mapping” studies, which are most meaningful when they differentiate between brain regions. Therefore, we tested whether the improvements found with MINDy-based Filtering are anatomically global or serve to further differentiate regions (i.e., are anatomically selective).

We consider two sorts of global effects: additive “shifts” in the global signal and global “scaling” of task effects. In statistical modeling terminology, the former reflects a main-effect (intercept) of method, whereas the latter reflects the method-specific slope. We modeled the differentiation between brain regions as either a main effect of regional significance (i.e., whether a region has a significant effect) or as an interaction with regional significance reflecting either a shift or rescaling of effect sizes of significant regions due to MINDy-based filtering, relative to the control models. We use the logical-valued variable *Sig_task,P arc_* to denote whether a parcel exhibited a significant effect for either method in a given second-level task analysis. We denote the MINDy-filtered second-level estimate (group-T) for each as *Y_task,P arc_* which is modeled as a function of matched control analyses (e.g. the original GLM or pre-regressed) which are denoted *X_task,P arc_*:

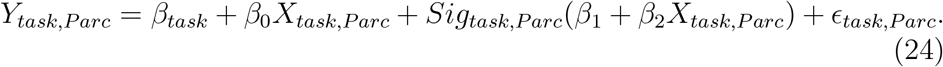

We assume that is independently and identically distributed across tasks and parcels (iid.). The coefficient *β*_1_ represents the main effect of parcel significance as a binary factor, while *β*_2_ represents the interaction with parcel effect size in control methods. Conceptually, these two components represent the degree to which MINDy-based Filtering further separates task-implicated and non-implicated parcels and the degree to which differences among task-implicated regions are further magnified, respectively.

Results indicate that the MINDy-based Filtering technique demonstrates differential sensitivity, in that improvements are greater in task-implicated regions (Fig. 6A). The main effect of event-related regional significance was significant relative both the original (*β*_1_ = .97 *± .*09; *t*(1669) = 10.8*, p ≈* 0) and pre-regressed pipelines (*β*_1_ = 1.05 *±.*09; *t*(1669) = 12.2*, p≈* 0). This result indicates that MINDy-based Filtering further separates event-implicated and non-implicated regions rather than simply increasing global statistical features. This feature also held at the single-task level in which linear models revealed a main effect of regional significance in all four tasks for both original (max *p* = .0007; Fig. 6B) and pre-regressed controls (max *p* = .0025). MINDy-based Filtering also differentially magnified effect sizes relative the original analysis (*β*_2_ = .075 *±.*023; *t*(1669) = 3.3*, p* = .001), but this effect was small and did not reach significance for the pre-regressed control (*β*_2_ = .034 *±.*022; *t* = 1.53*, p* = .13, 2-tailed) . Thus, task-implicated regions experienced the greatest improvements due to MINDy-based Filtering. For the current dataset, this approach primarily functioned to further highlight task-implicated brain regions (a main effect of regional significance) rather than magnifying the differences between task-implicated regions. These results imply that MINDy-based Filtering is sensitive to task-implicated brain regions rather than inducing anatomically global effects.

**Fig 6.**
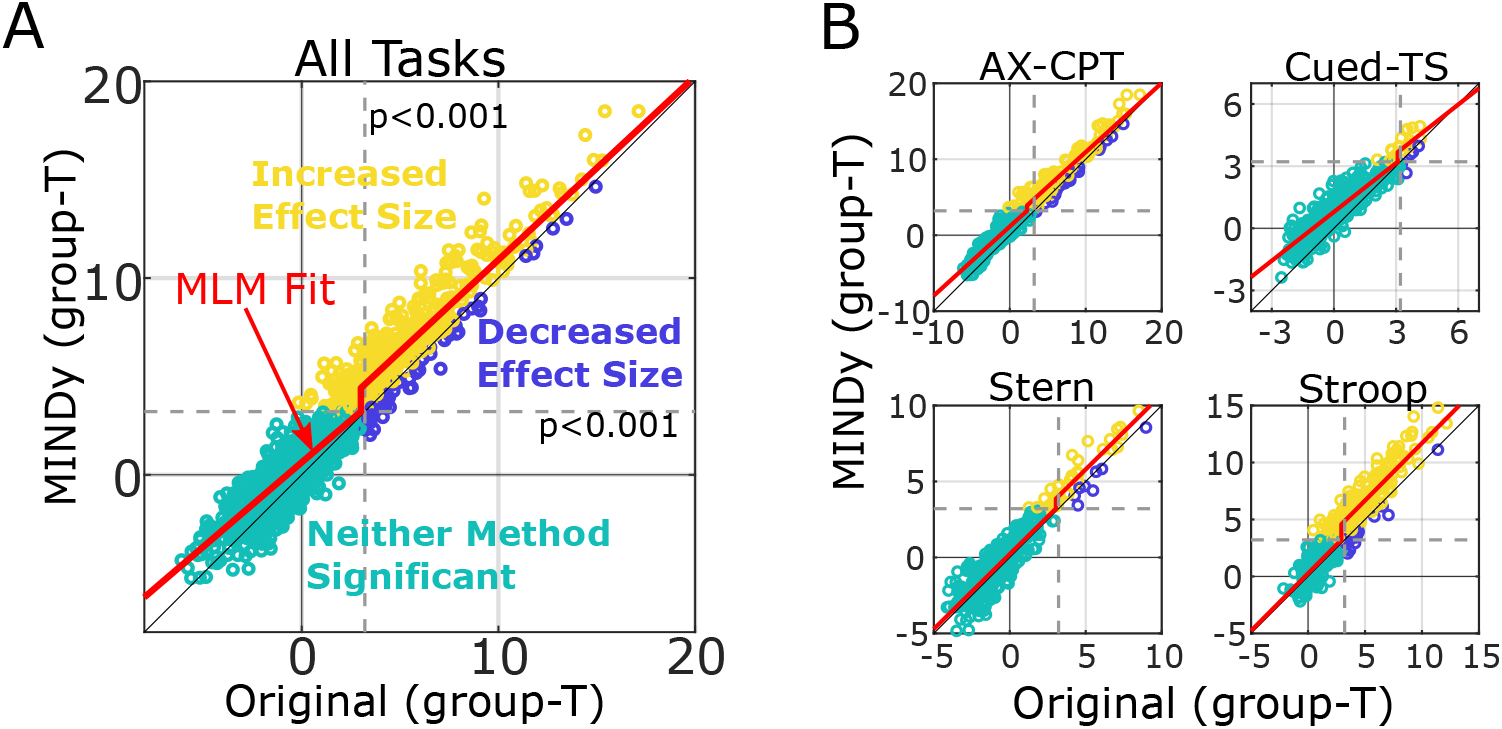
MINDy-based Filtering enhances task-related signals relative to controls. A) Comparison of parcel significance before and after MINDy-based Filtering collapsed across tasks. The multi-level model fit (averaged across the main effect of task) is plotted in red and the threshold-nonlinearity indicates sensitivity to parcel-significance. B) Task-specific comparisons relative the original analyses. Improvements can be seen in the number of parcels exhibiting higher t-values after MINDy-based Filtering relative to conventional analyses (i.e., above the identity line). Yellow dots indicate significant parcels (in terms of the control-demand effect) which also had increased effect sizes from MINDy-based Filtering, while blue dots denote significant parcels whose effect sizes were larger with conventional analyses. Teal dots denote parcels which did not exhibit a significant control-demand effect for either method. “MINDy” denotes MINDy-based Filtering and “Orig” denotes the original pipeline. Results with the pre-Regressed pipeline were indistinguishable from those with the original pipeline.

### 5.7. Identifying a Latent Cognitive Construct

The previous analyses indicate that MINDy-based Filtering enhances the identification of neural activity associated with a set of contrasts between trial-types (theoretical high control-demand trials minus low control-demand trials). However, many cognitive neuroscience studies seek to understand cognitive constructs, as opposed to unitary tasks. In the current section, we explore how well each method identifies the neural correlates of one such construct: cognitive control. The four tasks we studied have all been previously used to index cognitive control (typically via the difference between high-control and low-control trials). However, because the tasks themselves are not construct-pure (i.e., they tap multiple cognitive constructs) the neural activity associated with tasks is also expected to be non-identical. To control for this fact, we used the different trial types to generate levels of “construct-purity” in terms of cognitive control: low-control trials (low purity) and the high-vs.-low contrast (high purity). We consider the high-vs.-low contrast to be more “construct-pure” in terms of cognitive control since it controls for many of the other cognitive processes that differentiate tasks. For instance, speech production (unique to the Stroop task), is identical between high and low-conflict trials (the same set of words are produced). Likewise, working memory maintenance during delays (Sternberg, AX-CPT, and Cued-Task Switching) does not differ between high and low control-demand trials since these trial-types are identical through the delay period (up until the probe).

We tested how sensitive each approach was to the cognitive control construct via the relationship between “construct-purity” and cross-task similarity of neural effects. For this test, we indicate that a measure is “sensitive” to a factor (cognitive constructs) if the similarity in measurements reflects the similarity in that factor. We therefore consider a measure “sensitive” to cognitive constructs if it reports higher similarity between tasks for the high “construct-purity” condition (high-vs.-low control demand contrast) than for the low “construct-purity” condition (low demand trials).

We tested whether increasingly similar psychological contexts (conditions) across tasks are associated with more-similar neural effects using the Generalizability coefficient (a form of inter-class correlation/ICC; [40]). We compared measures in terms of their generalizability in tasks conditions which tapped a common construct (cognitive control demand) as well as conditions in which tasks were less psychologically similar. We predicted that MINDy-based Filtering would identify greater neural similarity between psychologically similar task conditions (higher generalizability/ICC) relative to psychologically dissimilar conditions, reflecting construct-selectivity. Conversely, we expect the ICC for psychologically disimilar task conditions (“low purity”) to be lower, reflecting disimilar neural activity patterns. The ICC “units of observation” consisted of the group-mean beta for each brain parcel (all 419 brain regions) and “classes” consisted of the different tasks. Results indicated that the proposed technique was sensitive to the cognitive control construct at group level (Fig. 7A). In the “low-purity” condition, with MINDy-based Filtering there was significantly lower similarity between tasks (*ICC* = .50 *±.*02) than the original and pre-regresed pipelines (*p^/^s < .*001, 5,000 bootstraps). Thus, MINDy-based Filtering does not generically increase the similarity of task results irrespective of cognitive construct. By contrast, for the “high-purity” condition, MINDy-based Filtering generated significantly more similar results across tasks (*ICC* = .60 *±.*03)than the original and pre-regressed pipelines (*p^/^s < .*001, 5000 paired bootstraps). We conclude that MINDy-based Filtering improves sensitivity to the cognitive control construct at group-level. Based on the nature of how these ICCs were calculated, the finding can also be interpreted as indicating that the anatomical profile of effects (i.e., the gradient of effect sizes across the brain) becomes more similar or consistent across tasks after MINDy-based filtering.

**Fig 7.**
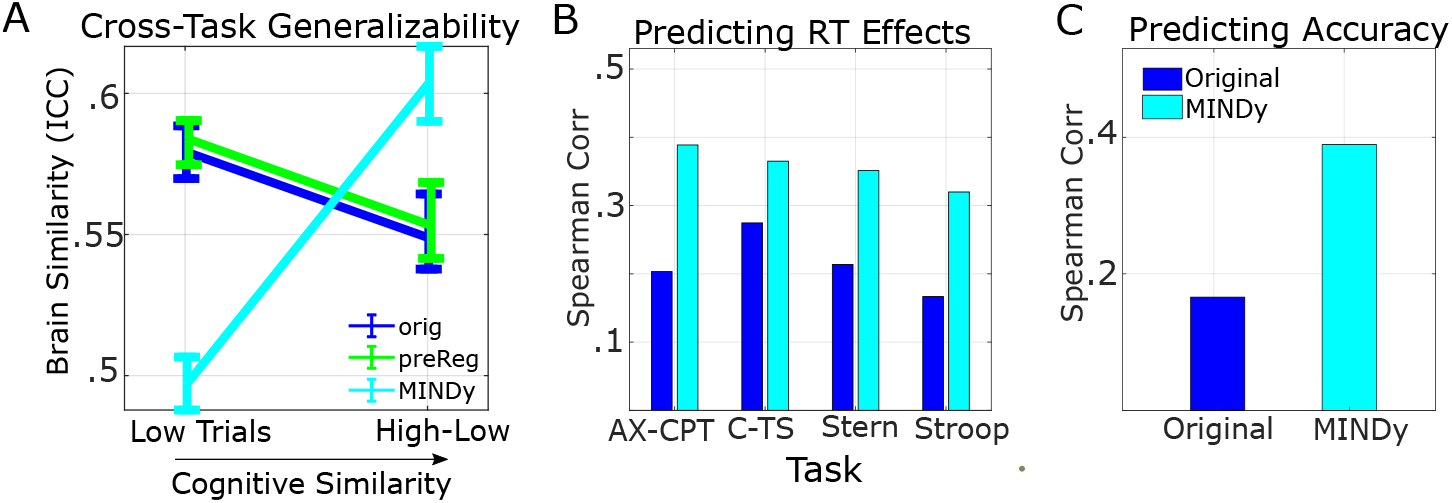
MINDy-based Filtering enhances cross-task similarity and behavioral prediction of cognitive control. A) MINDy increases the similarity of brain activation profiels (Generalizability Coefficient/ICC) between task conditions that engage similar psychological mechanisms (high-low cognitive control contrast) across tasks, but not when conditions do not isolate a common construct (low control trials). B) MINDy-based Filtering enhances correlations between event-related responses (average over DMCC34) and RTs in each task (collapsed across the 3 scanning sessions). C) MINDy-based Filtering also enhances the correlation between sustained responses in DMCC34 and error-rates (baseline session). Analogous results for the pre-regressed pipeline are displayed in SI. “MINDy” denotes MINDy-based Filtering, pre-Reg denotes the control pipeline with motion regression performed before GLM fitting, and “Original” denotes the conventional pipeline. Fig. 11C,D.

### 5.8. MINDy-based Filtering Enhances Brain-Behavior Relationships

The previous section demonstrated that neural effects identified with MINDy-based Filtering better generalized across task conditions tapping a common construct (cognitive control). In this section we demonstrate that this relationship also holds for behavior by using individual differences in task effects to predict the corresponding variation in behavioral cognitive control effects.

To isolate the effect of cognitive control demand we contrasted high-control and low-control trials for both the neural and behavioral data.

This approach, comparing trial types, is common in neuroscience including the neuroscience of individual differences. We found, using conventional analysis, that brain-behavior relationships were greater for the contrast between trial-types than for trial types in isolation. Averaged over tasks, the original pipeline had a mean correlation with RT of *ρ* = .21 for high-low vs. *ρ* = −.15 for high alone. The analogous correlations for MINDy were *ρ* = .36 (high-low) and *ρ* = -.08 (high only). For this reason, we employed the high-vs.-low control contrast in comparing methods.

For each subject task session, we summarized event-related effects in each task× method via the difference of normalized (z-scored over subjects) high and low control trial coefficients averaged over the DMCC34 set of parcels and similarly for sustained effects. Behavioral measures were similarly defined by the difference in normalized RT between high and low control trials and the accuracy in high control trials (nearly identical results are derived using high-low since low-trial accuracy is near ceiling). We found that MINDy-based Filtering increased the recorded correlations with RT for each task (Fig. 7B) and the average change in correlation across tasks was statistically significant (vs. original and vs. pre-regressed: *p < .*05, 5,000 bootstraps). Similarly, our approach increased correlations with accuracy (*p < .*05, 5,000 bootstraps, Fig. 7C). Interestingly, we found that across methods, individual differences in RT were positively correlated with the conflict-related (event) brain response but had a weaker relationship to sustained activity (Fig. 11A,B). By contrast, individual differences in accuracy were positively correlated with sustained activity, but unrelated to event-related activity (SI Fig 11A,B). Therefore, we compared methods in predicting RT using event-related estimates and in predicting accuracy using estimates of sustained activity. Results using the pre-regressed pipeline are depicted in SI Fig. 11 C, D. We conclude that after MINDy-based Filtering, individual differences in brain responses better predict behavioral measures associated with cognitive control.

## 6. Discussion

We demonstrated that MINDy-based Filtering increases the ability to detect both event-related (cognitive control-demand) and sustained brain responses in task fMRI (Sec. 5.5, SI Sec. 7.5). These effects are strongest in task-implicated brain regions (Sec. 5.6) and generate higher temporal precision than the original BOLD timeseries. By modeling and then partialing-out intrinsic dynamics, MINDy-based Filtering reduces both trial-to-trial variability within subjects, and variability between subjects (Sec. 5.4). However, while the absolute magnitude of subject-to-subject variability decreased, individual differences (and group—level activity) in a latent cognitive construct (control-demand) generalized better between tasks after MINDy-based Filtering (Sec. 5.7). MINDy-estimated task effects were also more predictive of individual differences in behavior (Sec. 5.8). These results suggest that MINDy-based Filtering can enhance the detection of task-evoked brain activity.

### 6.1. Relationship with Frequency-Based Filtering

Frequency-based (spectral) filtering has been applied to fMRI signals in many previous studies ([43],[44]). High-pass filtering is commonly applied to both resting-state and task data to remove signal drift which is thought to largely reflect changes in non-neuronal variables. Low-pass filtering is also sometimes applied, primarily for resting-state data. Although these approaches were common in early fMRI experiments, the changing nature of fMRI acquisitions (e.g. TR length) and analyses (e.g. functional connectivity) has led to renewed debate over these techniques ([45]) and the development of more sophisticated methodologies (e.g. [46],[47]). In the current work, we did not perform spectral filtering (instead using AFNI’s “polort” function for polynomial basis de-drifting). Likewise, MINDy-based Filtering is not a direct replacement for spectral filtering, which can be applied before our technique, afterwards, or not at all. However, as previously mentioned, when the connectivity parameters of the model are zero, the proposed technique reduces to a form of spectral filtering based purely upon autoregressive models. Empirically we have demonstrated that MINDy-based filtering outperforms filters based upon autoregressive models (SI Sec. 7.7, SI Fig. 10), so effects cannot be attributed solely to removal of particular frequency components within each region.

Notably, MINDy-based Filtering improves detection in both sustained and event-related analyses over both conventional methods and autoregressive filters. By contrast, filters based upon autoregressive models are expected to underperform in the identification of (low-frequency) sustained effects, as we confirmed in supplemental analyses (SI Sec. 7.5). At a statistical-level, dynamical systems models (including MINDy) capture the multivariate partial autocovariance between successive time-points (i.e. how *x_t_*_+1_ is related to *x_t_*). As a result, removing these predictions from the training data (Rest) inherently yields a timeseries with lower autocovariance. The improved detection of sustained effects is therefore significant as it indicates that MINDy-based Filtering reveals systematic differences between the resting-state and task dynamics rather than simply acting as a high-pass filter. These effects are also more pronounced in task-implicated parcels (Sec. 5.6, Fig. 6) indicating that these features are also context-related.

### 6.2. Relationship with other approaches

The current approach is conceptually related to several current initiatives for linking resting-state and task-state brain activity. Our approach uses resting-state brain dynamics to extrapolate patterns of intrinsic dynamics that also factor into brain activity during task states. Frameworks such as Activity Flow ([8]) have demonstrated similarity between the spatial aspects of evoked responses and resting-state network structure. Likewise, functional connectivity patterns have been found to be roughly similar between resting-state and task ([48]). However, whereas these frameworks are largely employed to discover similarities between spontaneous and evoked activity, we analyze the manner in which the task-state deviates from resting-state activity over short time-scales (how activity changes over short time-steps or TRs).

Other approaches have also investigated the difference between brain dynamics in task-state and resting-state. Previous work ([11], [10]) has demonstrated that intrinsic dynamics shape task-evoked activity on a trial-by-trial basis and modeling studies have reproduced the statistical differences between task and resting-state activity ([12]). Our approach furthers these efforts by leveraging these underlying concepts into an empirical modeling/analysis framework.

Dynamic Causal Modeling (DCM, [19]) frameworks have also used empirical dynamical systems models to improve estimates of task effects. As previously mentioned (Sec. 1.2), DCM techniques allow task effects to manifest changes in the exogeneous drive to brain regions and (for small-scale DCMs) the effective coupling between brain regions. By contrast, the current MINDy-based Filtering technique only models a single factor: changes in the input to each brain region, which collapses both of these mechanisms into a single term, as is also common in larger-scale DCM models (e.g. [21]). Our approach differs from all DCMs, however, in that we produce a timeseries of latent state estimate (task-related “input” to each region) which does not require any preconceived model of task effects (i.e., that they follow a certain temporal pattern). In the current work, we used statistical GLMs to analyze the MINDy-filtered data with Finite Impulse Response models fit for each trial type and additional components to model task blocks (mixed block/event-related designs). However, the end-product of our technique (a timeseries) could, in principle, be analyzed with a wide variety of methods, including parcel-level multivariate techniques (e.g., multivariate pattern analysis; MVPA).

### 6.3. Limitations

The proposed work rests upon three related claims: 1) intrinsic dynamics are roughly conserved between task periods and rest, 2) that by subtracting intrinsic dynamics we identify changes in “input” to each brain area and 3) that the signal generated by this calculation is a better marker of task effects (ostensibly task-related cognition). The first two claims are interdependent. We have mathematically defined changes in “input” as the signal components which are not explained by intrinsic dynamics (the residual after subtracting the modeled intrinsic component). The accuracy of estimated changes in “input” thus hinges upon whether the modeled intrinsic dynamics meaningfully generalize. We attempted to address this question empirically (see Sec. 5.3), and the results suggest that this assumption does hold. Specifically, we found that MINDy models estimate variation in the timeseries better during rest-blocks than task-blocks, which makes sense as the short rest blocks during task scans are more akin to resting-state scans. Moreover, even within task blocks, accuracy is well above chance and the timepoints that are not well explained by MINDy models (derived from resting-state) are precisely those during peak task effects (probe periods during each trial). Conversely, after MINDy-based Filtering these periods were well explained by task-based GLMs (with more variance explained than if MINDy-based Filtering were not applied) which indicates that the deviation from models is well explained by systematic, additive “input” to the model, as opposed to a breakdown in model-assumptions which would increase trial-to-trial variability. We also note that the generalizability assumption is “soft” in the sense that small changes in effective connectivity do not violate our assumptions. Since each connection describes the strength of input to the “post-synaptic” region, changes in connection strength are absorbed in the input estimate (summing over “pre-synaptic” sources). However, our assumption that MINDy-based Filtering removes mostly “nuisance variance” could be violated by some forms of large, systematic changes in effective connectivity. Fortunately, this assumption is easy to check (e.g., see Sec. 5.3) and we have not found evidence of its violation.

#### 6.3.1. Methodological Considerations

The bulk of our results concern the last claim (improved detection power) and the demonstration that observed statistical improvements are related to task-specific neural processes. We performed these tests using several controlled comparisons and lines of inquiry. However, our efforts in this domain are limited by using a specific subset of cognitive tasks: those used to index cognitive control. As the set of potential cognitive constructs remains vast, further testing in other cognitive domains may be useful.

Another limitation concerns how MINDy models are parameterized. Since we parameterize models based upon resting-state data, we require the collection of both resting-state and task data for each subject which increases data requirements. Moreover, this dependency could prove problematic for low-quality resting-state data, as mis-specified resting-state models could corrupt task estimates. We found that individual differences in goodness-of-fit were consistent across tasks (see SI Sec 7.2) so this possibility cannot be ruled out. However, previous analyses of MINDy modeling indicated that the goodness-of-fit is not related to individual differences in motion ([1]) and, similarly, MINDy-based Filtering was not impaced by individual differences in motion (SI Sec. 7.8). The results also do not support model overfitting, as goodness-of-fit did not decrease when applied to inter-block task periods (”rest” blocks) relative to training (rest) data (Fig. 3A). We also observed that using the group-average MINDy-Filter improved results relative conventional analyses (but less than individualized models; SI Sec. 7.3) so using a common MINDy Filter may ammeliorate short/low-quality resting-state data. Further study may therefore be beneficial in determining which factors (neural or nuisance) influence individual differences in goodness of fit, as these factors could influence estimated individual differences in task variables.

#### 6.3.2. Mechanistic Considerations

Future study is necessary is necessary to disambiguate which biological mechanisms contribute to the calculated “input” signal. For decades, computational neuroscience models have largely formalized task context as an exogeneous forcing (“input” or “bias”) term in neural networks and connectionist models (e.g. [49], [50], [51], [52], [53]). This formulation is appealing for its simplicity; however, external contexts serve only as “inputs” during sensory transduction, since brain activity is known to modulate even sensory neurons (e.g. [54], [55]). Even when these effects are neglected, many modeling studies assume that brain regions receive task “inputs”, even if these regions are not directly enervated by sensory nerves (e.g. [51]). As a result, these “inputs” should not be interpreted as literal inputs to the brain (i.e. signals from sensory nerves). Rather, these “inputs” include the initial propagation of such signals over the fMRI sampling rate (1 TR), so our approach is limited by the temporal resolution of fMRI BOLD.

The nature of these “inputs” is also somewhat underspecified. In the current approach, we use MINDy to model the propagation of brain signals during resting-state. The model predicts task-fMRI activation based upon the effective connectivity parameters estimated from resting-state. However, these parameters are limited to describing the relationship of bulk activity between brain regions. Many brain regions contain diffuse sets of neurons with heterogeneous axonal connectivity profiles. Several lines of evidence suggest that task-contexts can modulate the effective connectivity between brain regions via selective recruitment of neurons in synchronous ensembles ([56], [57], [58]). Our approach is therefore limited, in that it does not explicate how changes in “input” relate to changes in the effective coupling between brain regions. Future studies may improve upon the current approach by further modeling how task events modulate effective connectivity between brain regions. Such studies could either directly parameterize connectivity task interactions (as in DCM), or extend the filtering approach to estimate time-varying (or state-varying) connectivity.

### 6.4. Task Dynamics Could Potentially Influence Statistical Improvements

The current approach serves to estimate latent changes in input to each brain area. In the present study we found that MINDy-based Filtering consistently improved statistical detection power across tasks. However, there may be contexts in which brain activity (*x*(*t*)) is a more consistent marker of task context than input (*I*(*t*)). Such cases occur when different input patterns (i.e., inter-trial variability in input) lead to the similar outcomes in terms of activity. In these cases, MINDy-based Filtering might actually decrease detection power, since the “input” on each trial is less consistent than its long-term consequences. Future studies might identify such cases using a wider variety of tasks.

One area in which our approach could also be limited is in detecting slow neural events in which task-related activity evolves over multiple TRs. Since our approach acts as a pre-processing filter (i.e. doesn’t use task information) it is possible that it could filter out the propagation of very slow task-related activity in addition to task-unrelated activity. However, this cancellation is only expected when task-related activity propagates identically (has the same dynamics) to spontaneous brain activity. In practice, we have found that MINDy-based Filtering improves the detection of sustained brain activity and strengthens brain-behavior linkages (Sec. 5.8, SI Sec. 7.5).

### 6.5. Conclusion

In the current work, we proposed a new technique to estimate the influence of external contexts (task conditions) on brain activity (in our case fMRI). This technique forms a mathematical filter and therefore functions as a preprocessing step rather than as a direct tool for hypothesis testing. This property is advantageous as it allows this approach to be used in conjunction with a variety of existing methods. We have demonstrated that using MINDy-based Filtering improves statistical power (Fig. 5C), increases sensitivity to task-implicated regions (Sec. 5.6; Fig. 6)), and better identifies the neural signatures of a latent cognitive construct (cognitive conflict) (Fig. 7A). Moreover, MINDy-based Filtering enhances the strength of brain-behavior reslationships that differentiate subjects (Fig. 7B,C). These improvements are not sensitive to motion within a reasonable range (SI Sec. 7.8). Our technique can be easily inserted into most fMRI processing pipelines and we have made code available via the primary author’s GitHub to facilitate this process.

## 7. Supplemental Information

### 7.1. Relationship with Background-Activity

Our framework is conceptually related to that of background activity ([59, 60, 61, 62]) in which brain activity during task is modeled as the superposition of a canonical task-evoked response and trial-to-trial variability (“background activity”). In that approach, background activity is isolated by subtracting the task-related component as estimated during statistical GLM analyses, and it has been used to estimate Functional Connectivity during task ([59, 60, 61, 62]). However, despite both approaches dividing brain activity into two components, our approach fundamentally differs in terms of what signals are considered task-related vs. intrinsic. Nondynamic approaches divide the observed signal into systematic task effects and zero-mean “noise” (in the GLM sense) whereas dynamic frameworks consider both extrinsic and intrinsic contributions to how the brain evolves moment-to-moment. Passive downstream propagation of brain activity is predicted by intrinsic dynamics so these indirect effects are attributable to intrinsic factors despite being systematic (nonzero mean). As a result, these features remain in conventional GLMs but are removed during MINDy-based Filtering. We illustrate this point in a toy-model simulation featuring two linear nodes with a single directed connection and time-varying input to each node (Fig. 8 A,B). As the simulation indicates, MINDy-based Filtering extracts the timeseries of input to the system (Fig. 8 C) whereas downstream effects (i.e., the activation of n2 due to n1) are predicted based upon intrinsic dynamics (following the initial input; Fig. 8 D). By contrast, conventional GLM analyses do not separate direct and indirect processes and ascribe both features to the task-effect (Fig. 8 F, G). For this reason, the background activity and model predictions are not equivalent. Of course, unlike this toy simulation, neural processes occur over multiple timescales, many below fMRI resolution. As such, the estimated “input” actually reflects early processing and later active processing (as opposed to direct input from sensory nerves) and model-predictions reflect passive propagation of these signals over longer timescales. In our data, model predictions are more similar to the original timeseries than to the estimated “background activity”. Thus, although our approach has some conceptual relationships with the task-regression approaches to estimating background activity, these approaches are not equivalent and the intrinsic dynamics are not synonymous with background activity.

**Fig 8.**
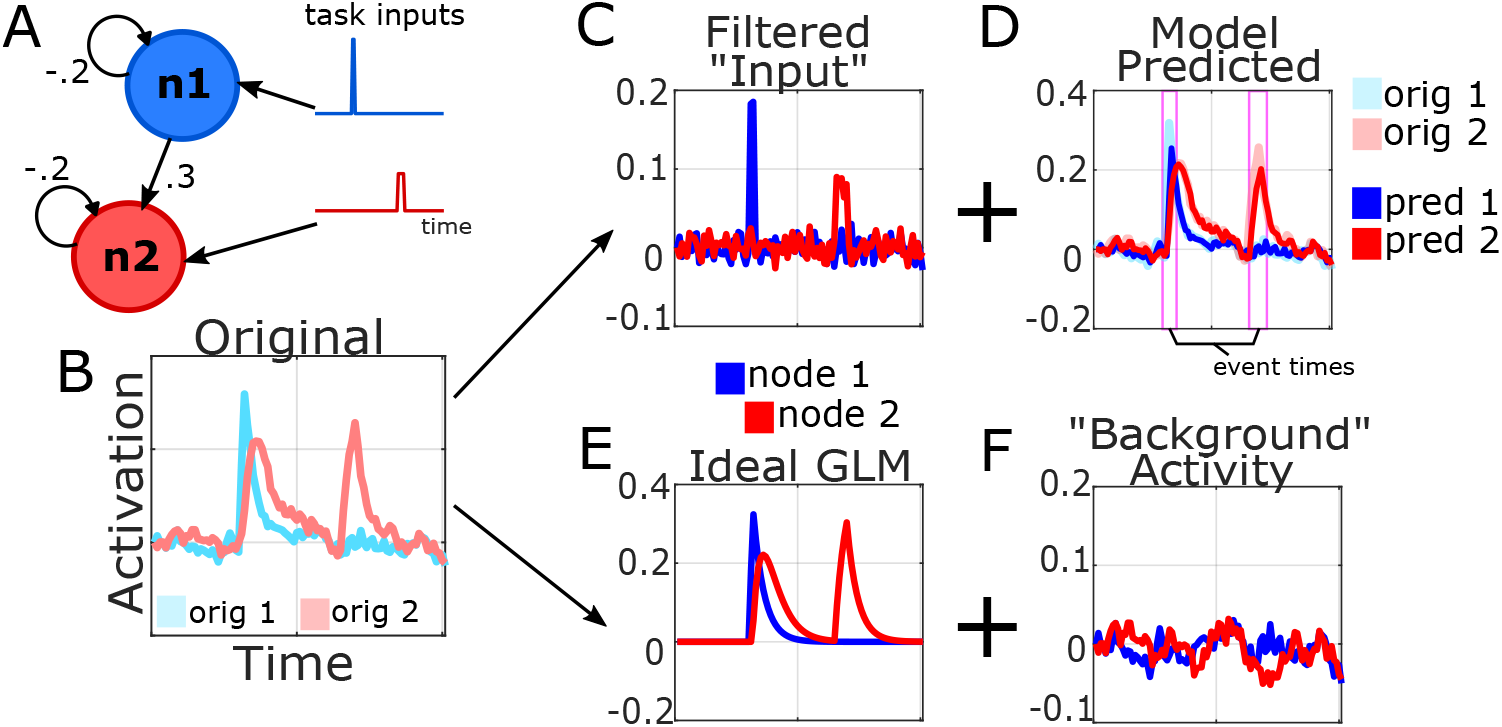
Comparison of signal decomposition via background-activity vs. MINDy-based Filtering. A) Toy model of a two node network with separate inputs to each node. B) Simulated timeseries. MINDy-based filtering decomposes the timeseries into the filtered “input” (C) and the model-predicted activity based upon intrinsic dynamics (D). By contrast, task-regression decomposes activity into a main-effect of task estimated by GLM (E) and “background activity” (F).

### 7.2. Sensitivity and Influences of MINDy Goodness-of-Fit

We found that model prediction accuracy was consistently lower for some subjects across all scan-types including the resting-state data to which the model was trained. This observation could reflect either model mis-estimation at rest or a general inability to predict that subject’s data even with a properly optimized MINDy model (due to poor signal quality or deviations from the MINDy framework). To distinguish between these possibilities we compared cross-subject prediction accuracy: the degree to which models trained to one subject’s resting-state predict another subject’s brain activity (rest or task). While cross-subject predictions were less accurate than within-subject (as expected), we found that the variation due to training-subject was far less than that due to testing-subject. Moreover, many subjects with poor model fits predicted other subject’s data better than their own. These results indicate that differences in model accuracy are primarily due to properties of poor-fitting subject’s data rather than the model fitting procedure per se.

We also tested whether our approach is sensitive to model goodness-of-fit. To test this influence we divided subjects into groups based upon median goodness-of-fit (either whole-brain or DMCC34 parcels) as measured during rest and during task (separately for each task). Analyses compared the mean T-value across parcels-of-interest for the two groups with pairwise parcels-of-interest defined as previously (at least one group passes *p < .*001 threshold). Null-distributions (10,000) were generated by randomly assigning subjects to two equal-sized groups without replacement. We did not find a significant difference in detection power for either task-combined data or any individual tasks using either resting-state or task goodness-of-fit (2 × 5 design). We conclude that improvements due to MINDy-based Filtering are not dependent upon goodness-of-fit within a reasonable range.

### 7.3. Influence of Individualized Brain Modeling

The primary restriction in applying our approach is the use of individualized brain models built from resting-state data. Acquiring sufficient data (we recommend ≥15 minutes) is time-consuming and may be particularly burdensome in special populations such as children. Therefore an important question for practical application is whether individualized brain models, as opposed to a single model, are necessary. This question is also theoretically interesting as it pertains to how individual differences emerge: via slow propagation along intrinsic dynamics or via the fast task “input” (dynamics below the fMRI TR). We address these questions in two sets of analyses.

In the first set of analyses we tested whether using a common MINDy filter, shared among subjects, is at least as powerful as individualized brain models. We defined a common MINDy filter by averaging the predictions of each subject’s MINDy model. We note that this procedure is not the same as using a common brain model as the parameters interact nonlinearly and covary. Hence the “average filter” cannot necesarily be inverted onto a single, representative MINDy brain model. Detection power using a common filter only slightly varied from using individualized models. For three tasks, the group-level filter performed significantly worse in detecting task events over the DMCC34 parcels (all but Cued-TS; max p=.01) .and for two tasks using the whole-brain (AX-CPT: *t*(175) = 8.39*, p* = 1.6*E* − 14; Stroop: *t*(244) = 5.49*, p* = 1.0*E* − 7) with differences in Cued-TS and Sternberg insiginificant. The combined detection power across tasks was significant for events (whole-brain: *t*(493) = 9.15*, p* = 1.6*E -*18; DMCC34: *t*(135) = 2.6*, p* = .01). However, individualized models only improved sustained effects over the DMCC34 parcel-set (*t*(135) = 3.52*, p* = .006) and not for the whole-brain analysis (*t*(293) = −1.1*, p* = .28). Interestingly, we also found little qualitative difference in terms of brain-behavior correlations (n.s.), suggesting that improvements reported in the main text are not dependent upon individual differences in resting-state.

We also repeated these analyses using random permutations of rest-subject and task-subject without replacement to test whether arbitrary assignments perform as well. Since our primary analyses concern group-level effects, using a group-average filter decreases noise and adds a further linkage between subjects. Using random pairings, as opposed to group-averages, thus provides a fairer comparison for identifying the influence of individual differences. Significance testing was performed using permutation tests (50,000 pairings of training/testing subject). As expected, random pairings performed worse than the group-average filter. We again found significantly worse detection power in event-related analyses compared to individualized models (average across tasks: *p < .*001; Cohen’s D=6.5; 50,000 permutations), but the absolute difference due to subject pairing was small (Δ*t* = .22 *± .*03) and the benefits over conventional analyses remained. We conclude that while individualized models do benefit power in detecting events, this effect is small relative the overall benefits of MINDy-based Filtering. We quantified the proportion of improvements due to individualized modeling as:

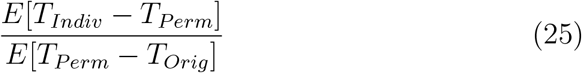

with *T_Indiv_* indicating the group-T of significant parcels for individualized MINDy and *T_perm_* indicating the corresponding values for random pairings of training (rest) and testing (task) subjects. Expectations are taken over task-relevent parcels (separately defined as in Sec. 5.2 for each permutation) and rest-task subject pairings. Results indicate that individualized models increased benefits in the cognitive-control effect by 32%, −1.1%, 16.5%, and 13.2% for AX-CPT, Cued-TS, Sternberg, and Stroop, respectively. The omnibus change (collapsed across tasks) was a 25.7% increase in benefits due to individualized models (i.e., most of the benefits for MINDy vs. orig remained). Thus, from a practial perspective, we believe that MINDy-based Filters constructed without individualized models can still significantly improve analyses above conventional methods, although further study is needed. Resultantly, the use of a single MINDy model (e.g., built from all subjects), as opposed to individualized models, may ease the requirements of quality resting-state data for each subject.

### 7.4. Influence of Deconvolution Parameter

We tested whether choice of the NSR (noise-signal-ratio) hyperparameter in Wiener deconvolution impacts results. This parameter dictates the degree of temporal filtering during deconvolution by regularizing the frequency-domain contributions. Larger NSR values leads to more filtering. We tested the influence of this parameter by repeating analyses with NSR chosen as .02, .005, .002 (main-text), or .0005. Thus, we tested NSR values ranging over a factor of 40. Results were highly similar for different values of the NSR parameters. Collapsing over subjects, parcels, and probe TRs, the high-low coefficient estimates correlated, on average, *r* = .99 across tasks and NSR combinations. Coefficients for the most dissimilar NSR parameters (.02 and .0005) correlated between *r* = .96 to *r* = .97 depending upon task. For comparison, the average correlation over tasks for MINDy vs. the original or pre-regressed pipelines was *r* = .73 and *r* = .71, respectively. All cases also preserved the benefits of MINDy-based Filtering. We conclude that, within a reasonable range, variations in choosing the Wiener NSR parameter do not strongly influence results.

### 7.5. Detection of Sustained Effects

MINDy also improved detection of sustained effects for the Sternberg and Stroop tasks relative the original and pre-regressed pipelines (max *p* = .0004; SI Fig. 9A). Trend-level improvements were observed in Cued-TS relative the pre-regressed pipeline (*t*(51) = 2.1*, p* = .04), but not relative the original pipeline (*t*(55) = 1.7*, p* = .10). However, sustained effects detected by MINDy did not differ relative the original or pre-regressed pipelines for the AX-CPT (*t*(103) = −1.1*, p* = .29*, t*(105) = −1.5*, p* = .14, respectively). Combined across tasks, MINDy increased detection of sustained events relative both the original (*t*(355) = 5.7*, p* = 2*E* − 8; SI Fig. 9B) and pre-regressed pipelines (*t*(353) = 6.2*, p* = 1.3*E -*9) as well as the autoregressive models (*t*(300) = 14.9*, p≈* 0*, t*(292) = 17.3*, p≈* 0 for global and local AR models, respectively; SI Fig. 10B). Thus, the proposed technique generally increased statistical power in detecting sustained effects. MINDy-based Filtering also increased the cross-task generalizability of group-average sustained effects (MINDy=.74 ± .04, all other pipelines *< .*65, *p < .*001, 5000 bootstraps). However, it’s important to note that sustained effects are not “construct-pure” and their distribution was highly skewed (strong visual component) so we urge caution in interpreting cross-task generalizability of sustained responses (although see Sec. 5.8 for its relevance to construct-specific behavior).

**Fig 9.**
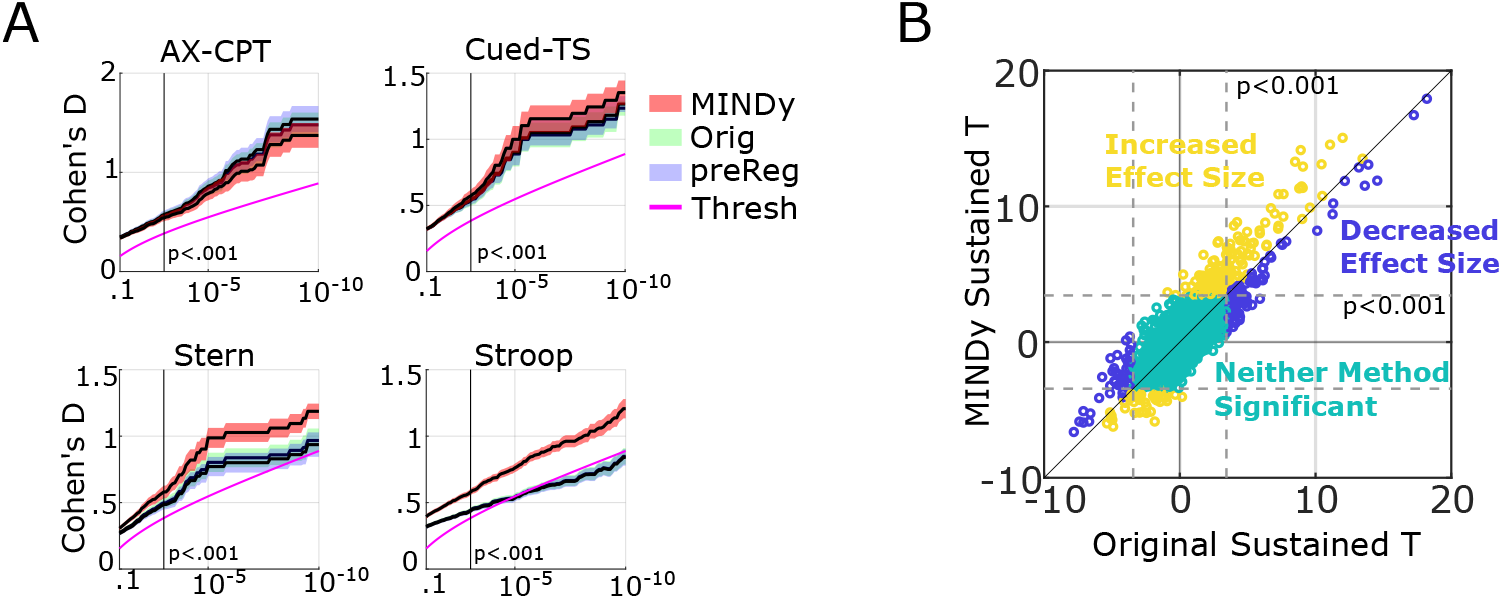
MINDy-based Filtering generally improves the detection of sustained effects. Unlike event-related effects, we permitted bidirectional sustained effects hence we compared the absolute magnitude of group-T statistics. The definition of significance was likewise 2-tailed. A) Pair-wise difference in detection power (group T) for the original pipeline and MINDy-based Filtering. B) Omnibus (task-collapsed) scatterplot of parcel significance using the original pipeline vs. MINDy-based Filtering for each task. Yellow dots indicate significant parcels (in terms of absolute sustained effect) which also had increased effect sizes from MINDy-based Filtering, while blue dots denote significant parcels whose effect sizes were larger with conventional analyses. Teal dots denote parcels which did not exhibit a significant control-demand effect for either method.

### 7.6. Sensitivity of Sustained Effects

As with event-related analyses, we examined whether improvements in the detection of sustained effects were limited to task-implicated regions. As before, we considered bidirectional effects for sustained analyses (i.e. parcels with significant increases or decreases in sustained activity). For this reason, we slightly modified Eq. 24 to model improvements in terms of magnitude rather than a linear main effect (*Y* again represents MINDy group-T, while *X* represents comparison pipeline group-T):

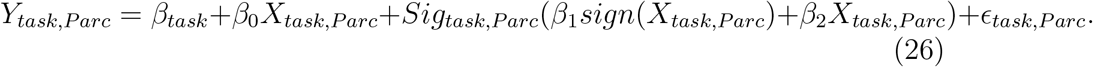

Note that the coefficient *β*_1_ is now multiplied *sign*(*Y_task,P arc_*). Results for sustained analysis mirrored those of the event-related analysis. As with event-related analyses, the proposed technique differentially increased effect sizes over task-implicated parcels when compared to the original, task-regressed, and global/local AR pipelines (*β*_1_ = .54*, .*68, 1.00, 1.03, respectively; max p=1.6E-8). As with event-related analysis, there was a slight trend of differential magnification vs. the original analysis (*β*_2_ = .05*, t*(1669) = 2.0*, p* = .048) but not vs. pre-regressed (*β*_2_ = .005*, t*(1669) = .22). We also observed a negative slope of *β*_2_, indicating diminishing returns (the opposite of differential magnification) relative the global (*β*_2_ = -.13*, t*(1669) = −5.4*, p* = 7.5*E-*8) and local (*β*_2_ =-.077*, t*(1669) = − 2.89*, p* = .0039) AR models. Thus, as with events, improvements under MINDy largely manifest a main-effect of parcel significance (i.e. increased categorical distinction between task-implicated and non-implicated parcels) as opposed to further differentiating among task-implicated parcels.

### 7.7. Comparison with Reduced Models

We compared estimation of inputs using MINDy models to analogous estimates to reduced autoregressive forms with autoregressive terms which were either subject-specific (but not parcel-specific) or terms which were specific to subject and parcel (see Methods Sec. 3.10). Since the MINDy model also features an autoregressive term (the “Decay”), these alternative models serve as reduced special cases which don’t include the effects of inter-regional signaling (connectivity). As such, improvements of the full MINDy model over these alternative (autoregressive) models indicate the contribution of modeling connectivity, as opposed to simply accounting for purely local dynamics.

Results indicated that group-level detection power for MINDy-based Filtering was greater than both the homogeneous and heterogeneous autoregressive comparison models. MINDy increased detection power over the both autoregressive models in terms of events over the DMCC34 parcels (max *p* = .0003) and for (whole brain) sustained effects (max p=2*E-6*; Fig. 9A). Whole brain analyses also indicated improved detection power for events in all tasks relative the global model (max p=.02) while all tasks other than Stroop (Stroop *t*(253) = 1.22*, p* = .22; other tasks: max *p* = 7.2*E* - 6) were improved relative the local model (SI Fig. 10A,B). There was a main effect of regional significance during multilevel modeling (i.e., improvement selectivity; see Sec. 5.6) for the proposed technique relative autoregressive comparison models (local: *t*(1669) = 3.87*, p* = 1.1*E* − 4, global: *t*(1669) = 4.15*, p* = 3.5*E* − 5). However, the proposed method did not significantly magnify effect sizes over AR pipelines (*p* = .16*, p* = .21 for global and local, respectively). Thus, the modeling of connectivity in MINDy primarily serves to further differentiate between task-implicated and non-implicated parcels as opposed to exacerbating differences among task-implicated parcels. MINDy-based filtering also improved the cross-task generalizability of cognitive-control effects relative autoregressive controls at both the group-level (local ICC=.50 *± .*03, global ICC=.52 *± .*03 vs. MINDy-based ICC=.60 *± .*03, *p < .*001, 5000 bootstraps).

**Fig 10.**
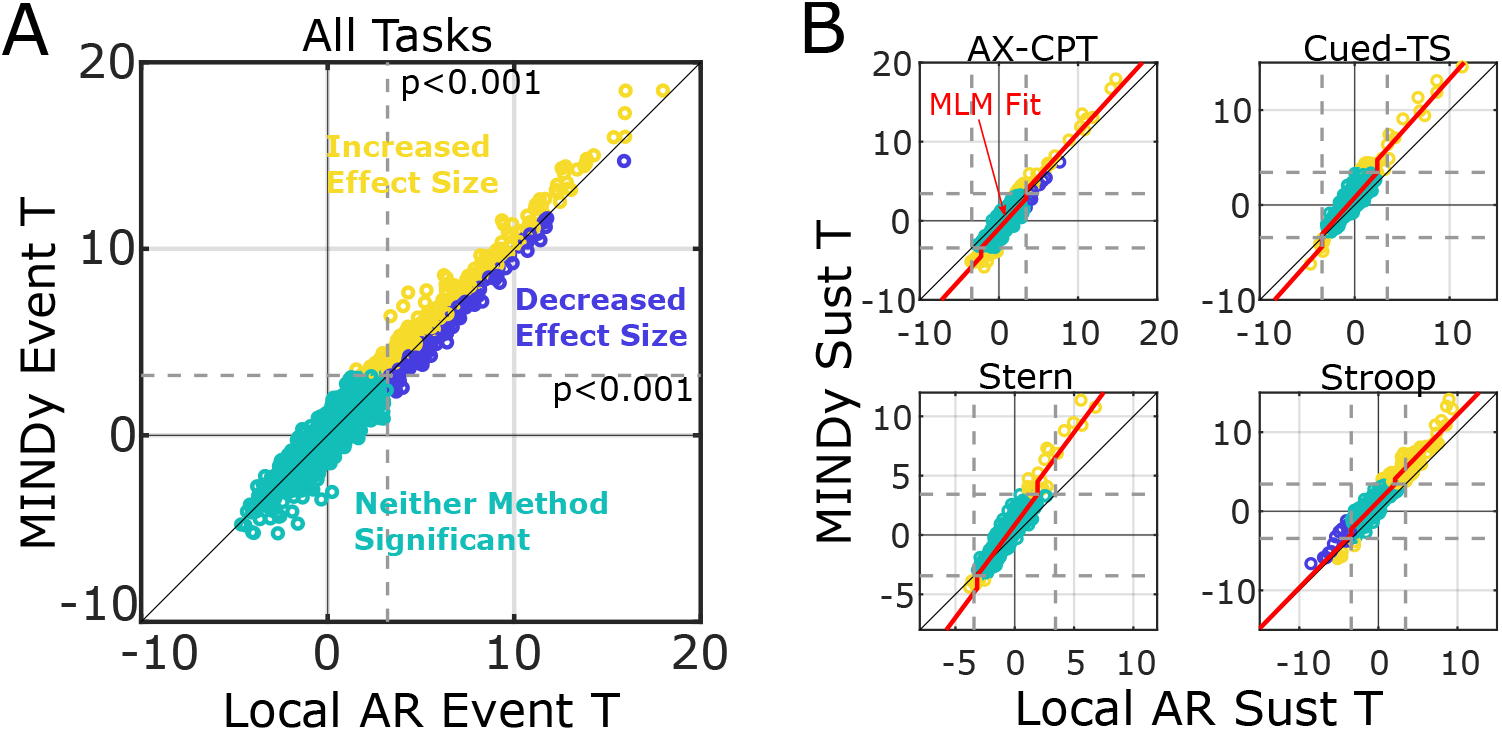
MINDy-based Filtering provides greater detection power using the full model over autoregressive (AR) reduced models which do not model connectivity. A) Omnibus (task-collapsed) scatterplot of parcel-wise event-related effects (high-low cognitive control demand contrast). Note that the improvements are smaller than those relative the original pipeline, indicating that some benefits in event-related detection are due to autoregressive filtering. B) Scatterplots of parcel-wise sustained effects when filtering with the local AR model vs. full MINDy model for each task. Note that AR pipelines perform worse than the original pipeline (larger MINDy improvement) for sustained effects. Yellow dots indicate significant parcels (in terms of the control-demand effect) which also had increased effect sizes from MINDy-based Filtering, while blue dots denote significant parcels whose effect sizes were larger with conventional analyses. Teal dots denote parcels which did not exhibit a significant control-demand effect for either method.

### 7.8. Sensitivity to Motion

Lastly, we compared the sensitivity of approaches to motion artifact. For each task and scanning session we computed three motion statistics: the number of frames censored due to passing a critical value of framewise displacement, the median framewise displacement and the median DVARS statistic ([41]) for each task run and averaged over runs. We then used resampling to test the relationship between each motion variable and the group effect-size of the high-vs.-low conflict contrast and sustained effect for each task. In brief, we randomly drew 5,000 samples of 30 subjects each without replacement. We computed group-level statistics for motion and the cognitive control contrast and then tested whether the average motion or variability of motion (inter-subject) of a sample predicted the sample’s group-effect (one-sample t-scores averaged over the 34 parcels). We also used the same technique for predicting the difference between methods (i.e. do improvements under our approach require low motion?). Results did not indicate a significant effect of motion for the current dataset and subject pool. The relationship between motion and the difference between methods (MINDy versus original averaged over tasks) was insignificant for event-related analyses and did not display a consistent sign (proportion of frames censored: *r* = .033, FD: *r* = −.078, DVARS: *r* = −.066). Likewise, we did not observe differential sensitivity to motion in the sustained effects (frames censored: *r* = .008, FD: *r* = −.011, DVARS: *r* = .01). Thus, the degree to which MINDy-based Filtering improves upon conventional methods is not influenced by motion within reasonable bounds.

**Fig 11.**
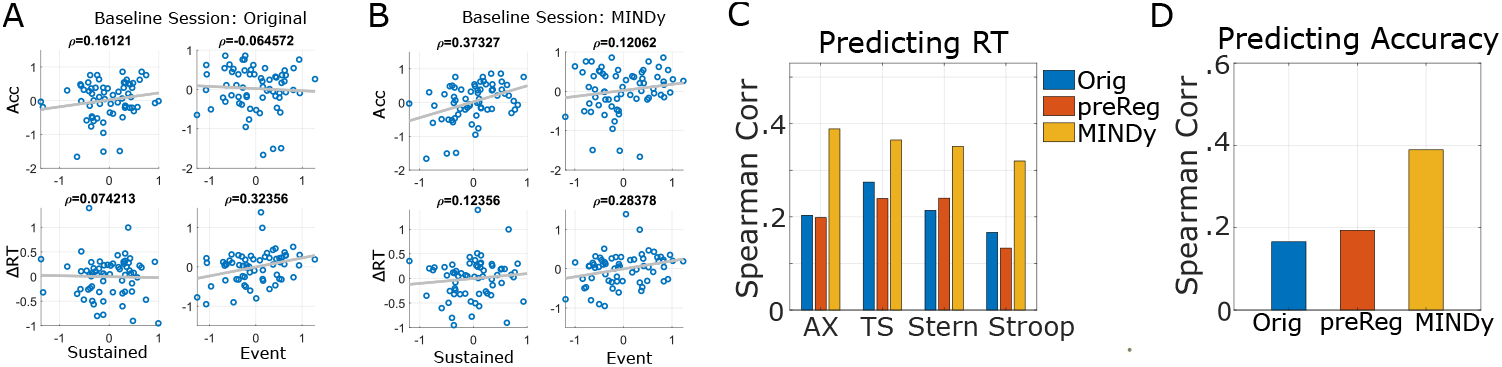
Predicting individual differences in behavior using brain activity (averaged over DMCC34). All measures are z-scored within task session. In the baseline session, individual differences in RT (averaged over task) are correlated with event-related brain activity, but not sustained activity for original and MINDy pipelines (A,B). By contrast, accuracy is predicted by sustained activity (B) but not by event-related activity (A,B). C) Individual differences in event-related activity better predict task RT (averaged over session) after MINDy-based Filtering relative the original and pre-regressed pipelines. D) Likewise, predictions of baseline-session accuracy using sustained activity (averaged over task) also increased. Panels C, D differ from the main text Fig. 7B,C by additionally including results for the pre-regressed pipeline.

## Acknowledgments

MS was funded by NSF-DGE-1143954 from the US National Science Foundation. TB acknowledges R37 MH066078 from the US National Institute of Health. SC holds a Career Award at the Scientific Interface from the Burroughs-Wellcome Fund. Portions of this work were supported by AFOSR 15RT0189, NSF ECCS 1509342 and NSF CMMI 1537015, NSF NCS-FO 1835209 and NIMH Administrative Supplement MH066078-15S1 from the US Air Force Office of Scientific Research, US National Science Foundation, and US National Institute of Mental Health, respectively.

